# Cortical temporal hierarchy is immature in middle childhood

**DOI:** 10.1101/678326

**Authors:** Dustin Moraczewski, Jazlyn Nketia, Elizabeth Redcay

**Affiliations:** Neuroscience and Cognitive Science Program, University of Maryland, College Park, MD; Computation and Mathematics for Biological Networks, University of Maryland, College Park, MD; Department of Psychology, University of Maryland, College Park, MD; Department of Cognitive, Linguistics, and Psychological Sciences, Brown University, RI

**Author notes:** Corresponding author. Address: Psychology Department, University of Maryland, College Park, MD, 20742, USA.

**Keywords:** development, inter-subject correlation, naturalistic viewing, social cognition, timescale

## Abstract

The development of successful social-cognitive abilities requires one to track, accumulate, and integrate knowledge of other people’s mental states across time. Regions of the brain differ in their temporal scale (i.e., a cortical temporal hierarchy) and those receptive to long temporal windows may facilitate social-cognitive abilities; however, the cortical development of long timescale processing remains to be investigated. The current study utilized naturalistic viewing to examine cortical development of long timescale processing and its relation to social-cognitive abilities in middle childhood – a time of expanding social spheres and increasing social-cognitive abilities. We found that, compared to adults, children exhibited reduced low-frequency power in the temporo-parietal junction (TPJ) and reduced specialization for long timescale processing within the TPJ and other regions broadly implicated in the default mode network and higher-order visual processing. Further, specialization for long timescales within the right dorsal medial prefrontal cortex became more ‘adult-like’ as a function of children’s comprehension of character mental states. These results suggest that cortical temporal hierarchy in middle childhood is immature and may be important for an accurate representation of complex naturalistic social stimuli during this age.

---

The cerebral cortex is organized along a hierarchy of multiple processing timescales (Hasson et al. 2008, Kiebel et al. 2008, Lerner et al. 2011, Honey et al. 2012). For example, primary sensory areas process transient incoming sensory information (i.e., short timescales) whereas regions higher in the temporal hierarchy reflect the integration and influence of information over multiple minutes or longer (i.e., long timescales) (Hasson et al. 2008). Using functional magnetic resonance imaging (fMRI), long timescale processing in the brain is typically indexed through examining synchronous neural activity between participants who have passively-viewed a naturalistic stimulus that has been scrambled at various resolutions (e.g., intact, coarse scrambled, fine scramble) (Hasson et al. 2008). While the functional role of cortical temporal hierarchy is beginning to emerge, no work has leveraged temporally manipulated naturalistic stimuli to explore the development of long timescale processing and its role in the development of higher-order social-cognition in children.

Previous work using naturalistic stimuli in adults suggests that cortical timescale processing reflects a process memory in which past information within a neural circuit affects the processing of newly arriving information (Hasson et al. 2015). Within this framework all neural circuits maintain the ability to accumulate information over time, though the specific time constant will vary depending on a given region’s location in the overall hierarchy (Lerner et al. 2011, Honey et al. 2012, Stephens et al. 2013, Hasson et al. 2015). The length of time that prior information can affect the processing of incoming information – known as temporal receptive window (TRW) (Hasson et al. 2008) – can be used to index a given region’s location in the temporal hierarchy. While the shortest cortical TRWs are seen in primary sensory areas, regions that display the longest TRWs are in higher-order association cortices and spatially overlap with the default mode network (DMN) (Hasson et al. 2010) – a network associated with internally directed thought, social cognition, and memory (Raichle et al. 2001, Buckner et al. 2008). In addition, regions of the DMN exhibit greater low-frequency power at rest and during natural listening relative to regions implicated in primary sensory processing, suggesting that regions within the DMN play a role in long intrinsic and extrinsic neural dynamics (Zou et al. 2008, Stephens et al. 2013). Further, activity within long timescale regions predicts scene recall during naturalistic viewing (Chen et al. 2016b, Simony et al. 2016, Chen et al. 2017), suggesting that these regions integrate information over multiple minutes and are important for following the long timescales of a story’s unfolding plot.

In addition to general memory processing, long timescales may be specifically important for real-world social-cognitive processing given that understanding the thoughts and intentions of other people requires the integration of information across multiple timescales (Zaki and Ochsner 2009, Hasson and Honey 2012, Koster-Hale and Saxe 2013). For example, to accurately assess what a social partner may be thinking during a conversation, one must integrate transient changes in their partner’s facial expression, word use, and prosody with the context of the current interaction, one’s previous knowledge of their partner’s personality, and one’s entire history of social norms and mores. In addition, while previous cortical temporal hierarchy work has suggested an overlap between the DMN and long timescale regions (Hasson et al. 2010, Chen et al. 2016b), other researchers have noted the overlap between the DMN and regions implicated in social-cognitive processing (Spreng et al. 2009, Mars et al. 2012, Li et al. 2014). This involvement of DMN regions associated with long timescales may be due to the fact that social-cognitive processing relies on integration across long time windows. Disruption in a narrative for example will lead to difficulties in tracking a character’s mental state. Indeed most prior work investigating long timescales has used movies or narratives rich with social content. However, the role of long timescale processing in higher-order social-cognitive processing remains to be directly investigated.

Middle childhood (between 6- and 12-years old) is an ideal time to investigate the aforementioned relationship between long timescales and social cognition. Within this age window, social-cognitive abilities increase (Apperly et al. 2011, Rice et al. 2016) and brain regions supporting social-cognition become more specialized during tasks (Saxe et al. 2009, Gweon et al. 2012, Warnell et al. 2017) and at rest (Fair et al. 2009, Supekar et al. 2010, de Bie et al. 2012, Muetzel et al. 2016). For example, the temporoparietal junction (TPJ) exhibits selectivity for mental state representation (Saxe and Kanwisher 2003), which increases with age throughout middle childhood (Saxe et al. 2009, Gweon et al. 2012). Further, children’s social-cognitive abilities predict the selectivity of the right TPJ for mental state representation (Gweon et al. 2012). At rest, the DMN exhibits greater within- (Supekar et al. 2010, de Bie et al. 2012, Muetzel et al. 2016) and between-network connectivity (Fair et al. 2009) with age. Thus, regions of the social brain (including the DMN) are functionally immature during middle childhood. One possibility is that maturation of long timescale processing within these regions is a domain-general mechanism supporting advances in social-cognitive development; however, no study has investigated long timescale processing in middle childhood.

Much of the previous developmental work is ill-equipped to address questions of long timescale processing due to the use of non-naturalistic stimuli that focuses on either discrete events (i.e., task-based) or no task (i.e., resting state). However, the use of naturalistic stimuli with fMRI to answer developmental questions is growing (Cantlon and Li 2013, Emerson et al. 2015, Moraczewski et al. 2018, Richardson 2018, Richardson et al. 2018) due to its similarity to real-world processing (Zaki and Ochsner 2009, Hasson and Honey 2012) while minimizing task demands and head motion (Vanderwal et al. 2015, Vanderwal et al. 2018). In a previous naturalistic viewing study, we demonstrated that adults exhibited greater neural synchrony in regions topographically similar to the DMN compared to 4- to 6-year old children and that responses in the left TPJ within the child group became more ‘adult-like’ as a function of age (Moraczewski et al. 2018). Other recent work suggests that functional maturity within the theory of mind network, as well as the pain network, is positively related to anticorrelation between the two networks (Richardson 2018, Richardson et al. 2018). Taken together, this work demonstrates that regions associated with the DMN are also functionally immature during naturalistic viewing; however, no work has specifically investigated timescale processing and its relation to cognition in childhood.

The current study examines developmental differences in long timescale processing and its relationship to social-cognitive abilities in middle childhood. Functional MRI data were collected on a cross-sectional sample of children and adults while watching two episodes of a children’s television show. To establish brain regions implicated in long timescale processing, each participant viewed one episode intact and one episode that was scrambled to induce temporal inconsistency. We first examined low-frequency power during the intact episode through sampling regions at different locations within the temporal hierarchy (Stephens et al. 2013). Further, to ground our analysis within the social-cognitive literature, we conducted our analysis within regions implicated in social-cognitive processing (e.g., dmPFC, TPJ, precuneus) (Schurz et al. 2014). Next, we utilized inter-subject correlation (ISC) analysis to examine differences in neural synchrony between the child and adult groups, as well as child-to-adult ISC as a measure of individual differences in the child group (also known as neural maturity (Cantlon and Li 2013)). To examine ISC at the group level, we used novel crossed random effects analysis, which has been previously shown to control the false positive rate more adequately than previous group ISC methods (Chen et al. 2017). In addition, to index social-cognitive processing, after the scanning session we assessed participants’ comprehension of the character’s mental states (mental comprehension) and events that required general memory of the episode but did not require knowledge of character mental states (non-mental comprehension).

We hypothesized that regions previously implicated in long timescale processing (e.g., those topographically similar to the DMN) would show increased low-frequency power compared to a control region in the primary auditory cortex. Further, we hypothesized that adults would exhibit greater low-frequency power relative to the child group. We hypothesized that regions of the DMN would show greater neural synchrony during long timescale processing (i.e. the intact episode) in the adult group and that, due to cognitive and neural immaturity, children would display reduced specialization for long timescales in these regions. We also hypothesized that long timescales would be more important for mental state comprehension compared to non-mental comprehension. Finally, we hypothesized that child-to-adult ISC in long timescale regions would be related to child age and comprehension. The current study is the first to map age-related differences in neural dynamics and long timescale processing, as well as relate long timescale processing to individual differences in higher-order social-cognitive abilities.

## Methods

### Participants

A cross-sectional sample of fifty-two children (30 female, 9.20 ± 1.95 years old, age range: 6.08–13.08) and twenty-seven adults (16 female, 22.76 ± 2.26 years old, age range: 19.25–26.75) were recruited to participate in the study. All participants were screened to ensure that they were native English speakers and had normal or corrected-to-normal vision. In addition, inclusion criteria required no history of neurological or psychological disorders, and no first-degree relatives with autism or schizophrenia based on self-report or parent-report for children. Families were recruited through a university database where parents opt-in to be contacted for research studies and received financial compensation as well as a toy for their time. Adult participants were recruited from the local undergraduate student body and received either course credit or financial compensation. After exclusion based on failure to complete all four functional MRI runs (4 children), failure to pay attention to the stimuli (e.g., eyes closed for an extended period of time) (1 adult), hardware malfunction (1 child and 1 adult), previous familiarity with the stimuli (1 child), and excessive motion (15 children and 1 adult, see Methods), the final sample consisted of thirty-one children (19 female, 9.83 ± 2.02 years old, age range: 6.42 – 13.08) and twenty-four adults (14 female, 22.08 ± 2.29 years old, age range: 19.25 – 26.75). All protocols were approved by the University of Maryland Institutional Review Board and implemented accordingly.

### Stimuli

Functional data were acquired while participants viewed two episodes from the first season of the Australian ‘mockumentary’-style children’s television show Little Lunch (Butler and Hope 2015). This show was chosen because 1) based on piloting, it was unlikely that children in the population from which we recruit would have familiarity with the show, 2) the show’s characters are of similar age to the population of interest (i.e., middle childhood), and 3) the mockumentary-style enables viewers to receive the mental state information from multiple characters regarding a single plot event. The Monkey Bars (MB) episode consisted of two girls arguing over whose turn it is to play on the monkey bars and The Body Bus (BB) episode consisted of a girl who thinks she has head lice and is worried that a visiting vehicle may be a doctor who is doing checkups. Prior to the scan session all participants were asked if they had previous experience with the show. Only one child participant was familiar and subsequently excluded from the study (see Participants). The beginning and end of each episode was cropped to the same length (11 minutes and 12 seconds). The length of time for the episodes was chosen so that each episode could be cropped to be the same length while preserving all meaningful plot content.

Each participant viewed both episodes: one episode intact and the other episode scrambled according to the follow procedure. To generate the scrambled version of each episode, we determined timestamps that corresponded to naturally occurring camera cuts or, if a scene was longer than 20 seconds, a naturally occurring break in the dialog. We chose a maximum scene length of 20 seconds to maintain consistency with previous work that investigates mid-level timescale processing (Honey et al. 2012, Chen et al. 2016b). In addition, no sentences were split to ensure that the full dialog was preserved in both the intact and scrambled versions of each episode. No statistical difference in scene duration between the two episodes was detected (t(168) = 0.43, p=0.67; The Monkey Bars – scene duration: 7.98 ± 4.06 seconds, range: 1.79 – 19.82 seconds; The Body Bus – scene duration 7.71 ± 4.02 seconds, range: 1.64 – 19.27 seconds). Once the scene cuts were determined, we pseudo-randomized the scene order to ensure that each scrambled episode was as disjunct as possible. In addition, in order to control for lower-level visual and/or auditory effects of cutting and reassembling the scrambled video, we cut and reassembled the intact version of each episode using the same procedure with a chronological rather than a random scene order. Finally, we divided each of the four episodes (i.e., MB intact and scrambled; BB intact and scrambled) in half (e.g., MB intact 1^st^ half and 2^nd^ half) in order to distribute each episode across two functional runs, which yielded eight stimulus files. Dividing each clip over two shorter functional runs helped to maximize participant compliance. The video files were edited using the Moviepy python module (https://zulko.github.io/moviepy/). All participants saw each of the episodes but one was presented intact and one scrambled (and this ordering was counterbalanced across participants).

In the scanner, participants passively-viewed an intact episode and scrambled episode which were each divided into two runs for four functional runs total per participant: intact run 1 (1^st^ half of intact episode), intact run 2 (2^nd^ half of intact episode), scrambled run 1 (1^st^ half of scrambled episode), and scrambled run 2 (2^nd^ half of scrambled episode)). The specific scrambled episode (i.e., MB or BB) and the sequence of presentation for the intact and scrambled episodes (i.e., intact then scrambled, or scrambled then intact) were counterbalanced. However, participants always viewed the 2^nd^ half of each episode (scrambled or intact) in the run immediately following the presentation of the first half of a given episode. Order A consisted of BB (“Body Bus”) intact and MB (“Monkey Bars”) scrambled whereas order B consisted of MB intact and BB scrambled. Sequence one consisted of intact then scrambled presentation whereas sequence two consisted of scrambled then intact presentation.

Our final sample size consisted of 11 adults and 15 children who viewed order A, 13 adults and 16 children who viewed order B, 12 adults and 15 children who viewed sequence one, and 12 adults and 16 children who viewed sequence two. Child age did not differ between order A and B (t(29)=0.73, p=0.47) or sequence one and two (t(29)=0.39, p=0.70). Each video was preceded with 10 seconds of fixation on a black screen and followed with 20 seconds of post-stimulus fixation. Videos were presented using Psychopy presentation software (version 1.83.04) (Peirce 2007), projected to a screen, and viewed though a mirror mounted to the head coil. Sound was presented using Sensimetrics S14 insert earphones.

### Episode Comprehension

Immediately after the scan, participants were given an assessment to index episode comprehension that required the knowledge of characters’ mental states, as well as comprehension that did not require mental state knowledge. Sixteen comprehension questions – eight mental and eight non-mental – were created for each episode (i.e., MB and BB episodes). To create the questions, we ran a behavioral pilot with 36 questions per episode (18 mental and non-mental). The final 16 comprehension questions were then chosen based on those that elicited the greatest individual response variability (see Appendix 1 in the Supplementary Information for final comprehension questions). Episode questions were presented in the same order in which participants viewed the episodes in the scanner. Prior to the questions, participants viewed a series of sixteen screen shots for each episode in order to disambiguate the specific episode to which the questions corresponded. Importantly, none of the screen shots contained the answers to any of the subsequent comprehension questions. Each screen shot appeared on the screen for 3 seconds followed by a 0.5 second fixation cross. Immediately after the screen shots, free-response questions appeared in text on the bottom of the screen and the researcher read the question aloud. Participants were also provided with images of the six main characters and their names. The comprehension assessment was presented on a Macbook Pro using Psychopy (version 1.83.04) (Peirce 2007).

Successful performance on the mental questions required the participant to understand the beliefs, desires, and/or intentions of one or more of the characters (e.g., “Why did Debra Jo ask the kids to calculate their own change?”), whereas successful performance on the non-mental questions did not (e.g., “What sport were Atticus and Rory playing?”). Answers to each question were scored as 1 for correct, 0.5 for partly correct, and 0 for incorrect or “I don’t know” (see Appendix 1 in the Supplementary Information for the scoring scheme we used for each question). These scores were then converted to percent correct for the mental and non-mental questions separately yielding four comprehension scores per participant (intact/scrambled x mental/non-mental). We used a linear mixed-effects model to estimate the effects of condition (intact/scrambled), question type (mental/non-mental), condition by type interaction, and age on the child participant’s comprehension scores, while controlling for episode (i.e., MB or BB). In addition, to account for repeated measures, a random effect of participant was added. The analysis of comprehension in the adult sample was analogous to the child comprehension except that we removed the effect of participant age as a predictor. After we estimated the effects of condition, question type, condition by type interaction, and age (in the case of the child comprehension), the confidence intervals for the uncertainty of the estimates were examined using a simulation procedure based on the effect mean and standard error (Gelman and Hill 2007). Our model results are reported as the fixed effect estimates and the confidence intervals surrounding their uncertainty. Finally, we also conducted pairwise post-hoc t-tests on the factors of condition and question type and corrected for multiple comparisons using a Bonferroni correction.

### Image Acquisition

Structural and functional MRI images were acquired on a Siemens 3T MAGNETOM Trio scanner using a 32-channel head coil. Each session began with four functional runs of echo planar images (EPI) (intact run 1, intact run 2, scrambled run 1, and scrambled run 2) followed by a high-resolution structural T1-weighted anatomical scan. For each functional run, 293 volumes were acquired using a multiband slice sequence, which consisted of 66 slices per volume, a voxel size of 2 x 2 x 2.2 mm, TR of 1250 msec, TE of 39.4 msec, and 90 degree flip angle. The first four functional volumes of each run were automatically dropped to allow for magnetization equalization. The structural T1-weighted MPRAGE image consisted of 192 contiguous sagittal slices, with a voxel size of 0.9 mm isotropic, TR of 1900 msec, TE of 2.32 msec, inversion time of 900 msec, and a flip angle of 9 degrees.

### Image Preprocessing

Structural images were processed using Freesurfer’s (version 5.1.0) automated segmentation and cortical surface reconstruction algorithm (*recon-all*) (Fischl 2012). This algorithm uses voxel intensity to assign each voxel to a tissue type (e.g., cortical gray matter, white matter, lateral ventricle). The white matter and pial surfaces are then used to construct a two-dimensional surface mesh representation of the participant’s brain. Two independent trained research assistants then checked the automated algorithm for accuracy. If necessary, edits were made to the cortical and white matter surfaces and the algorithm was repeated.

We utilized a surface-based analysis to preprocess the functional images using the AFNI (Cox 1996) and SUMA (Saad and Reynolds 2012) software packages (Version 17.2.13). We chose a surface-based processing pipeline to minimize possible bias in normalizing children and adult brains to a common 3D stereotaxic space (Jo et al. 2007). Preprocessing steps included slice-time correction followed by volume co-registration to ensure that each participant’s functional, structural, and surface datasets were registered within the participant’s original space. The white matter and pial surfaces created from Freesurfer were then used as a mask to project the functional data to a standardized surface mesh (36,002 nodes per hemisphere) using a mean-mapping function. We then normalized the BOLD signal to a mean of 100 through scaling the time series by the average node-wise intensity. The data were then entered into a nuisance regression, which included terms for linear, quadratic, and cubic low-frequency drift, de-meaned motion parameters and their derivatives, and average signal from tissues of no interest (i.e., white matter and lateral ventricle, defined from Freesurfer). In addition, we censored time points where the frame-to-frame displacement was greater than 0.5mm in any translation or rotation (Power et al. 2014). The residuals from the nuisance regression were smoothed using a 5mm FWHM Gaussian kernel on the surface. Finally, we concatenated the time series from each half episode (e.g., intact run1 and intact run 2) into one time series per episode. To do this we dropped the first 13 and the last 10 time points within each run (i.e., first 16.25 and last 12.5 seconds, respectively) and concatenated runs one and two of a given episode. The first thirteen time points were dropped to remove the pre-episode fixation and onset of the stimulus, while the last 12.5 seconds accounts for the tail end of the post-episode fixation. We maintain the first 7.5 post-episode fixation seconds to account for hemodynamic lag. Once the temporally cropped data were concatenated, all participants had two runs of preprocessed BOLD time series: one corresponding to the intact and one to the scrambled episodes.

To ensure that our results cannot be attributed to head motion, we then used the motion parameters generated during volume co-registration to further exclude participants. Since our planned analysis required each participant to contribute all four functional runs, our criteria for participant exclusion was met if *any* of the four functional runs exceeded 10% of time points with greater than 0.5 mm frame wise displacement (FD). This motion exclusion yielded 38 participants in the child group and 24 in the adult group. However, after our a priori motion exclusion, we detected a statistically significant difference in mean FD between the Child and Adult groups (t(60)=2.69, p<0.01), such that the Child group exhibited greater motion than the Adult group. Thus, we then quantified outliers in mean FD using a median absolute deviation greater than 2 (Leys et al. 2013). This analysis yielded seven motion outliers in the Child group (two 6-year olds, four 7-year olds, and one 9-year old). After the exclusion of the motion outliers, our final sample consisted of 31 participants in the Child group and 24 participants in the Adult group, with no statistical differences in mean FD detected between group (β_group_ = 0.014, 95% CI [-0.007, 0.034], t(62)=1.30, p=0.20). However, in the Child group, we did observe a negative and statistically significant effect of age on mean FD. Thus, all of our analyses that examine the age-related differences include child head motion as a covariate (see Supplementary information for group- and age-related analysis of head motion).

### Regions of interest

For both our low-frequency power and inter-subject correlation analyses, we extracted values from four regions of interest (ROIs) (Figure 1a). Three ROIs (temporoparietal junction (TPJ), precuneus, and dorsal medial prefrontal cortex (dmPFC)) were chosen to sample regions previously implicated in social cognition, the DMN, and long timescale processing. These regions were defined using coordinates from a meta-analysis of theory of mind tasks (Schurz et al. 2014). Bilateral ROIs were defined for the TPJ (MNI coordinates: −53, −59, 20 and 56, −56, 18), whereas one region was defined for each the dmPFC (MNI: −1 54 24) and precuneus (MNI: −4 −52 30). In addition, to examine the specificity of our effects, we chose bilateral control ROIs in the primary auditory cortex to represent a region that we did not predict to be involved in social cognition or long timescale processing (MNI: −50, −24, 3 and 54, −24, 0). Auditory regions were defined using the meta-analytic tool Neurosynth (Yarkoni et al. 2011) using the peak coordinates from the map associated with the search term ‘sound’. Data from the bilateral regions (i.e., TPJ and auditory) were averaged across both hemispheres for all subsequent ROI analysis and visualization (see Supplementary Information for analysis of separated TPJ hemisphere regions).

**Figure 1.**
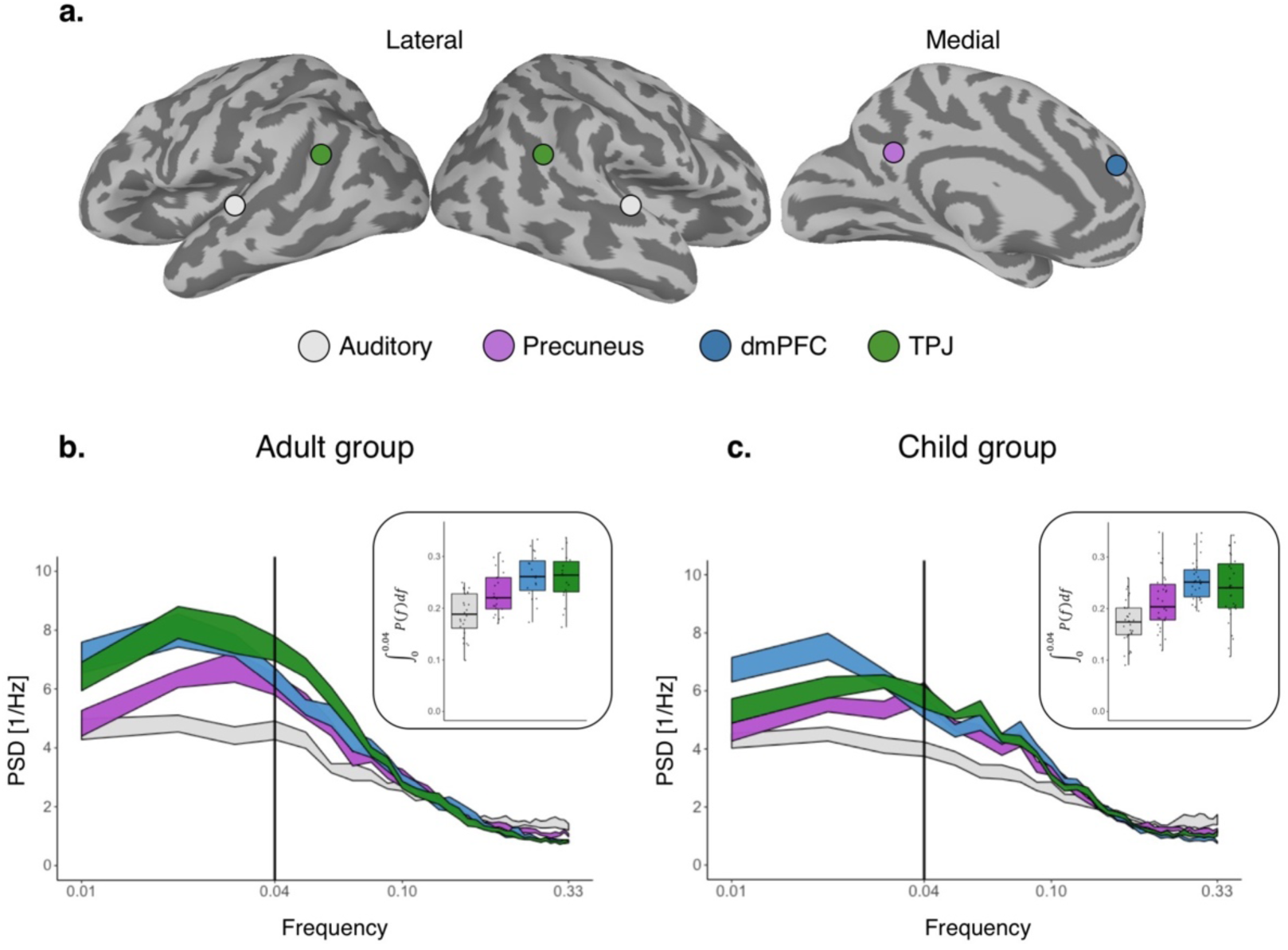
Low-frequency neural dynamics. (a) Regions of interest and power spectrum density (PSD) for the adult (b) and child (c) groups. The PSD ribbons for each region depict the mean ± standard error. The black line denotes the 0.04 Hz cut-off used in the low-frequency statistical analysis. The boxplots denote the proportion of low-frequency power for each region.

### Low-frequency Power Analysis

To characterize the neural dynamics of long timescale processing, we examined the proportion of low-frequency power within a priori ROIs while participants viewed the intact episode. Similar to the methods used by Stephens and colleagues, for each participant we began by removing the global mean from each node’s time series (Stephens et al. 2013). We then estimated the power spectrum for each node on the surface using the Welch method with a window of 100 seconds and 50% overlap. The node-wise power spectra were then averaged across all participants within the adult and child groups and the mean spectra were extracted from each ROI for further analysis. Using the overall frequency range of 0.01 Hz to 0.33 Hz, we quantified the proportion of low-frequency power within the 0.01 to 0.04 Hz band. Finally, we used a mean bootstrap procedure that randomly sampled participants within-group to examine the confidence intervals surrounding the mean power spectrum density (Stephens et al. 2013).

### Inter-subject Correlation

To examine neural synchrony within and between conditions and groups, we constructed pairwise inter-subject correlations (ISCs) for each surface node. Here we only calculated meaningful ISCs, that is, correlations between time series from participants who viewed the same episode of the same condition. Since roughly half of the sample saw the BB episode scrambled whereas the other half viewed the same episode intact, ISCs were only calculated within participants who viewed the same episode order (e.g., Order A or B, see Participants). The ISC for each pair of participants within the same order was calculated as the Fischer’s z-transformed Pearson’s product-moment correlation between the time series of each pair of participants. Node-wise correlations were calculated using AFNI’s *3dTcorrelate* function and normalized using *3dcalc*. Pairwise ISCs were calculated (as opposed to a leave-one-out average group procedure (Lerner et al. 2011)) in order to 1) utilize a crossed random effects analysis (Chen et al. 2017) and 2) examine individual differences in child-to-adult ISCs (Cantlon and Li 2013, Moraczewski et al. 2018).

### Statistical Analysis of Inter-Subject Correlations

Parametric statistical modeling of ISC data necessarily violates the assumption of observation independence, such that each participant’s data is represented *N - 1* times in the model matrix. Thus, our statistical analysis utilized fully crossed random effects in order to accurately estimate the shared variance between each effect (Chen et al. 2017). This method has been shown to more adequately control the false discovery rate compared to previous non-parametric methods of within- and between-group ISC statistical analysis (Chen et al. 2016a, Chen et al. 2017). For each episode and condition, we constructed a symmetric pairwise ISC matrix for each node on the surface where each row and column corresponded to one participant. Excluding the diagonal (i.e., self-self correlations), all pairwise ISCs were entered into the model (i.e., both bottom and top triangles of the matrix). Such symmetrical redundancy allows for an accurate estimate of the shared variance of each effect (Chen et al. 2017). In addition to fixed effects of interest, a random effect was added *for each* participant contributing to the corresponding outcome ISC value. Thus, we predicted each outcome ISC from a combination of fixed effects of interest and two random effects that correspond to each of the two participants contributing time series data to the outcome correlation.

Similar to previous work (Hasson et al. 2008, Lerner et al. 2011, Chen et al. 2016b), we examined regions implicated in long timescale processing by first calculating significant within-condition ISCs (i.e., Intact and Scrambled) for each group (i.e., Child and Adult). Here our effect of interest is the intercept (corresponding to the group mean ISC), while controlling for a fixed effect of episode (since data from both episodes are represented within each condition) and fully crossed random effects. Using this procedure, we calculated regions that show statistically significant ISC for each episode and condition, yielding four ISC maps: adult intact, adult scrambled, child intact, and child scrambled. To determine regions implicated in long timescale processing, we entered the within-group ISC data for both conditions into another mixed-effect model. Here the effect of interest was condition while controlling for episode and crossed random effects. This analysis yielded one intact – scrambled map for each group. To examine group differences in the within-group intact – scrambled maps, we entered all ISC data into another mixed-effect model. Here our effect of interest was a group x condition interaction, while also controlling for episode and crossed random effects. Finally, while we did not have a priori hypotheses regarding the effect of episode content, we controlled for this effect in all aforementioned analyses. See Supplementary Figure 5 for content-specific differences (MB versus BB) in long timescale specialization.

All whole-brain maps are presented using a node wise threshold of p < 0.01 and a cluster extent of 112 mm^2^ to threshold the data and maintain a FWE of p < 0.05. We calculated the cluster extent threshold using a Monte Carlo simulation (1000 iterations) in which we generated a volume of noise for each participant, projected it to the surface using a mean mapping function, and calculated cluster sizes that could arise by chance. Note that current cluster-based methods to control FWE have not been evaluated using crossed random effect or ISC analysis. In addition, we extracted ISCs from each of the four a priori ROIs (auditory, dmPFC, precuneus, and TPJ) for further analysis. All whole-brain and ROI models were built using the lmer() function in R. All analysis code is available at the following repository: https://github.com/dmoracze/TRW.

### Individual Differences in Child-to-Adult Inter-Subject Correlations

We next examined cortical regions in the Child group that become more ‘adult-like’ as a function of individual differences in age and social-cognitive comprehension. To index social-cognitive comprehension, we calculated a composite score of mental / (mental + non-mental) episode comprehension to examine regions that show greater ISCs as a function of mental state comprehension beyond general episode comprehension. Rather than using within-group ISC (i.e., adult to adult or child to child correlations), we used the child-to-adult ISCs (Moraczewski et al. 2018), a metric also known as neural maturity (Cantlon and Li 2013). For this analysis, we predicted child-to-adult ISC as a function of the fixed effects of a covariate (i.e., age and mental comprehension composite score), condition, and covariate by condition interaction, while also controlling for differences between episodes and crossed random effects. In addition, for both child-to-adult ISC models we controlled for child head motion and, for the mental state composite model, we also controlled for child age. Since our questions revolved around long timescale processing, our main effect of interest in the child-to-adult models was the covariate by condition interaction, such that we hypothesized that age and mental state comprehension would be more related to child-to-adult ISC in long timescale regions within the Intact compared to the Scrambled episode. All maps are presented using a node wise threshold of p < 0.01 and a cluster extent of 112 mm^2^ to threshold the data and maintain a FWE of p < 0.05.

## Results

### Children exhibit less low-frequency power in long timescale regions compared to adults

We extracted the mean power spectrum density from the four a priori ROIs (auditory, dmPFC, precuneus, and TPJ) and quantified the proportion of low-frequency power (0.01-0.04 Hz) within each ROI. Consistent with previous work (Stephens et al. 2013), we found a statistically significant difference in low-frequency power between the four ROIs in the adult group (F(3,69)=25.56, p<0.0001), ensuring that we controlled for head motion (using mean FD) and accounting for four within-participant repeated measures. Follow-up tests suggest that the dmPFC, precuneus, and TPJ all exhibited greater low-frequency power compared to our control auditory region (dmpfc: β_region_=0.10, 95% CI [0.08, 0.13], t(23)=8.83, p<0.0001; tpj: β_region_=0.11, 95% CI [0.08,0.14], t(23)=7.62, p<0.0001; precuneus: β_region_=0.05, 95% CI [0.03,0.07], t(23)=5.08, p<0.0001 – reported effects are in units of proportion of low-frequency power (0.01-0.04Hz) relative to the entire PSD frequency range (0.01-0.33Hz), all p values are Bonferroni corrected) (Figure 1b). Further, controlling for head motion, the child group exhibited a similar pattern of neural dynamics compared to the adult group, such that we detected a difference in the proportion of low-frequency power between regions (F(3,90)=22.23, p<0.0001) and the dmPFC, precuneus, and TPJ exhibited greater power compared to the auditory region (dmpfc: β_region_=0.09, 95% CI [0.07,0.11], t(30)=8.49, p<0.0001; tpj: β_region_=0.07, 95% CI [0.05,0.09], t(30)=6.44, p<0.0001; precuneus: β_region_=0.04, 95% CI [0.02,0.06], t(30)=3.84, p<0.001) (Figure 1c). In comparing the low-frequency power between groups and controlling for head motion, we found that the adult group exhibited greater proportion of low-frequency power compared to the child group in only the TPJ (tpj: β_group_=0.06, 95% CI [0.02,0.10], t(52)=2.85, p<0.05; dmpfc: β_group_=0.03, 95% CI [0.00,0.06], t(52)=1.88, p=0.26; precuneus: β_group_=0.03, 95% CI [0.00,0.06], t(52)=1.69, p=0.39; auditory: β_group_=0.01, 95% CI [-0.01,0.04], t(52)=1.23, p=0.89, all p values are Bonferroni corrected) (Figure 2). Finally, in the child group, we examined the relationship between the proportion of low-frequency power and child age. Controlling for head motion and repeated measures, we did not observe a main effect of child age on proportion of low-frequency power (F(1,28)=1.23, p=0.28). We did observe a trend within the dmPFC such that older children exhibit greater proportion of low-frequency power, however this result did not survive the correction for multiple comparisons (tpj: β_age_=0.00, 95% CI [-0.01,0.01], t(28)=0.01, p=1.00; dmpfc: β_age_=0.01, 95% CI [0.00,0.02], t(28)=2.12, p=0.17; precuneus: β_age_=0.01, 95% CI [-0.01,0.02], t(28)=0.94, p=1.00; auditory: β_age_=0.00, 95% CI [-0.01,0.01], t(28)=0.35, p=1.00, all p values are Bonferroni corrected) (Figure 3). Notably, for the purposes of the aforementioned analysis, we combined the left and right TPJ regions into one region, however, our results remain consistent when examine the left and right TPJ regions individuals (see Supplementary Figure 2).

**Figure 2.**
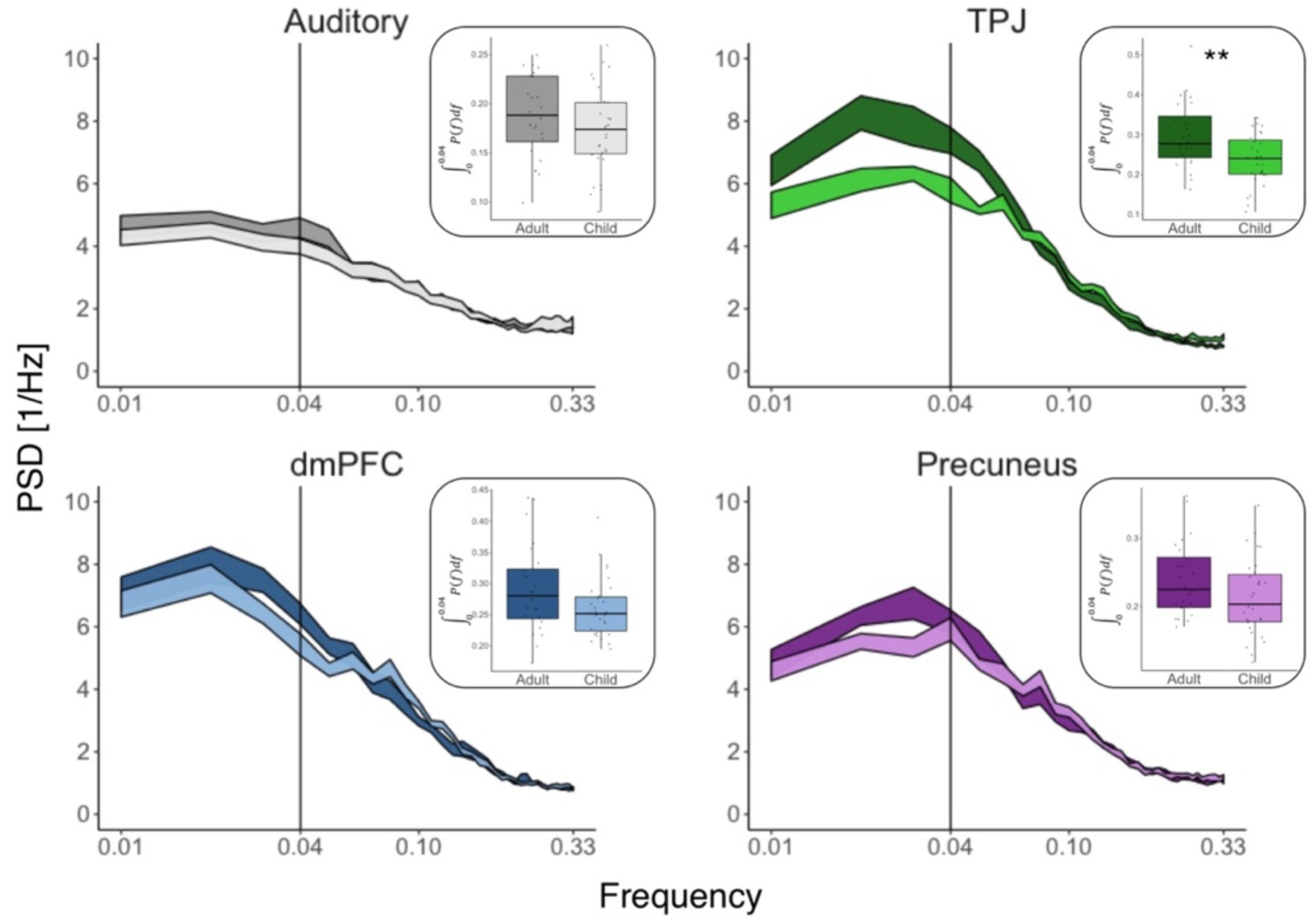
Differences in low-frequency neural dynamics between child and adult groups. The power spectrum density (PSD) and proportion of low-frequency power are presented for each group and region. The adult group is shown in dark colors while the child group is in lighter colors. The PSD ribbons for each region depict the mean ± standard error. The black line denotes the 0.04 Hz cut-off used in the low-frequency statistical analysis and the boxplots denote the proportion of low-frequency power for each region.

**Figure 3.**
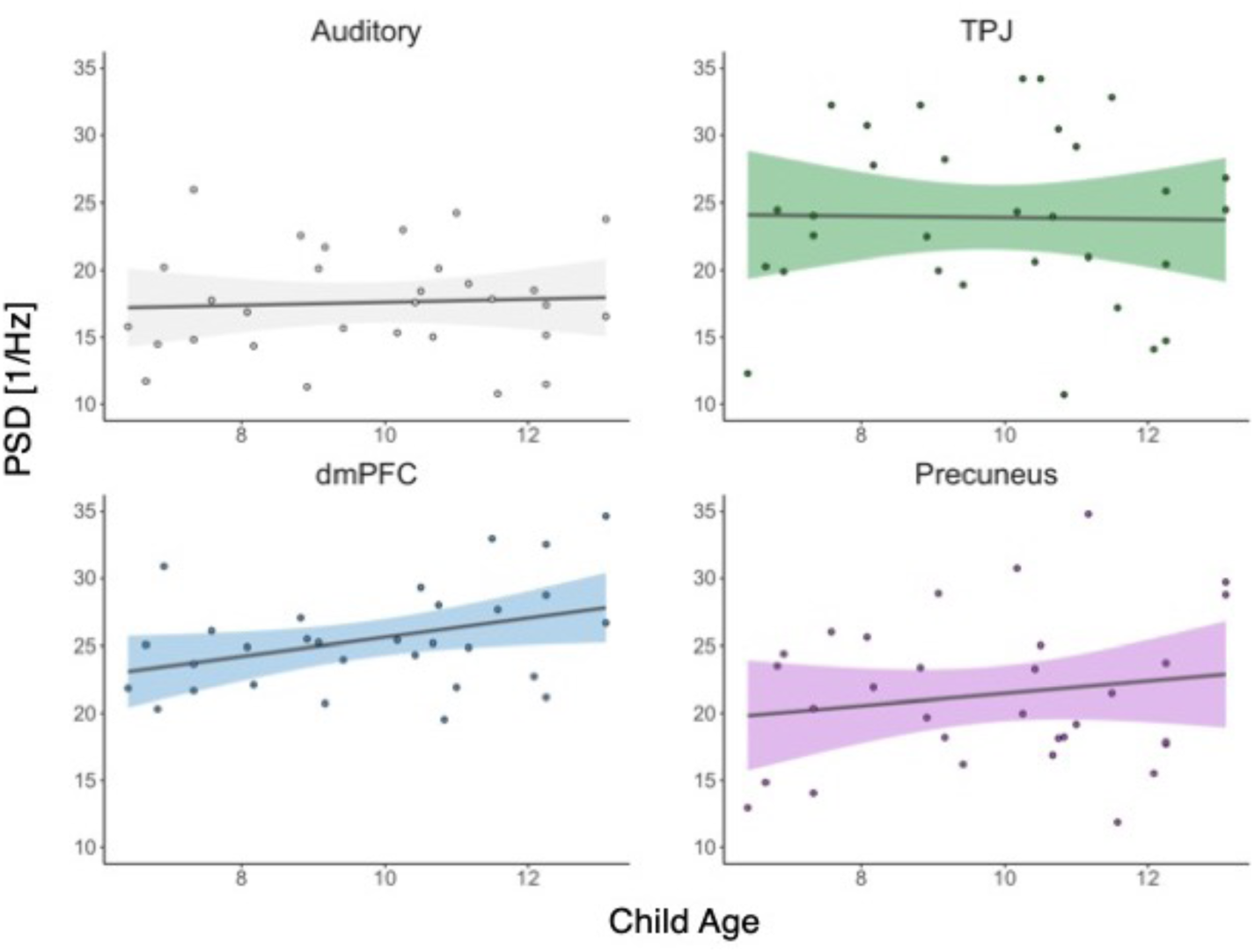
Age-related differences in low-frequency power within children. For each region, we plot the relationship between the proportion of low-frequency power and child age. A regression line is imposed onto the scatterplot with the shared areas denoting the 95% confidence interval on the beta estimate and colored by region.

### Children exhibit less neural synchrony for long timescales

To examine the influence of long timescales on within-group ISC, we first constructed whole brain within-group models for each condition and group separately. Here the effect of interest was the intercept (i.e., group ISC mean) while controlling for differences between episode and crossed random effects. We found that both the Intact and Scrambled conditions elicited statistically significant ISC in primary and secondary visual cortices, superior temporal gyri, superior temporal sulci, inferior temporal cortex, posterior and dorsal parietal areas in both the Adult and Child groups (nodewise p<0.01, cluster extent 112 mm^2^, FWE p < 0.05) (Figure 3a,b, respectively).

**Figure 3.**
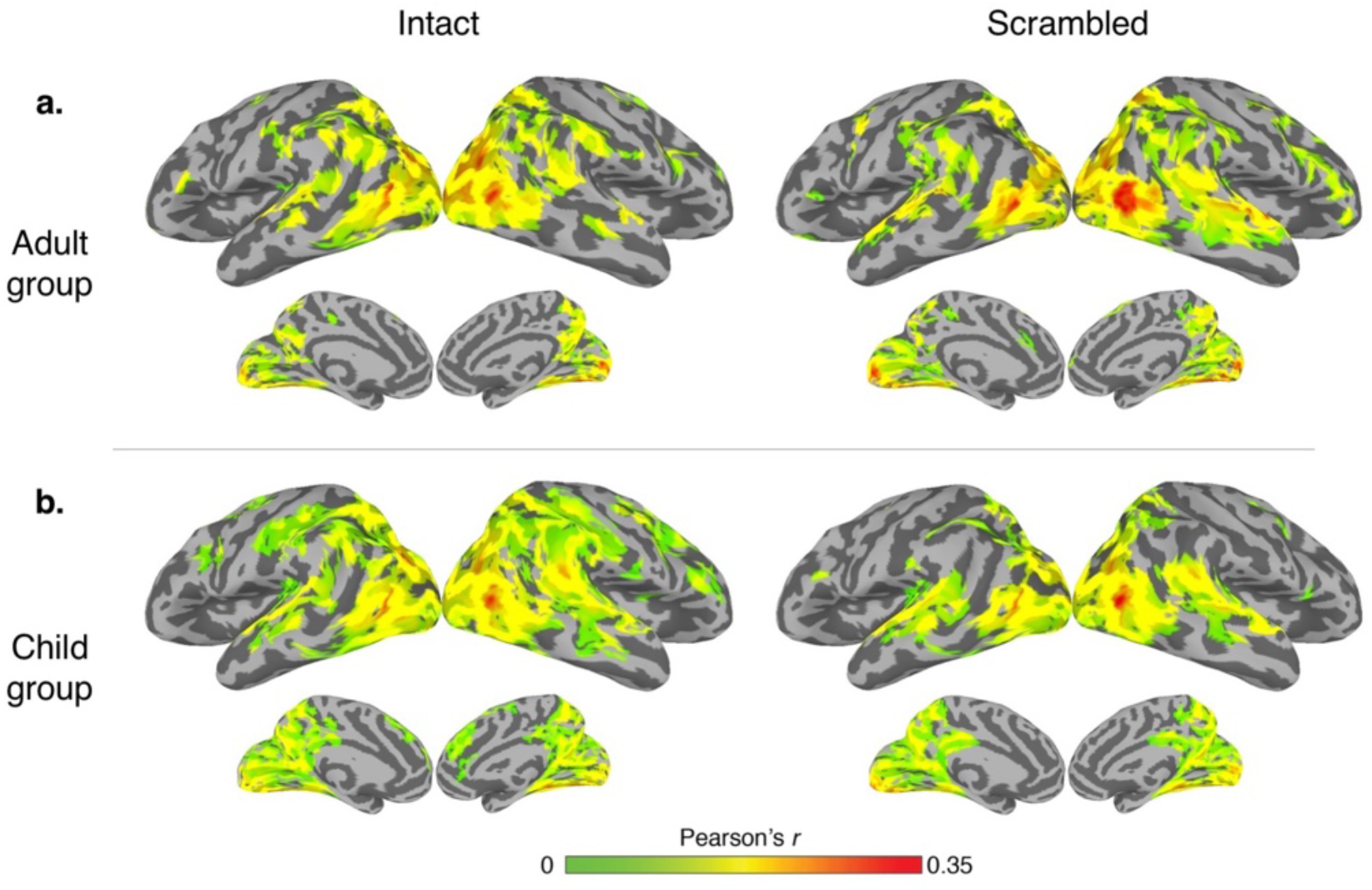
*Within-group inter-subject correlation*. Within-group ISC maps for the adult (a) and child (b) groups. Maps are thresholded with a nodewise p < 0.01 with a cluster extent of 112 mm^2^ to achieve a FWE of p < 0.05.

We next investigated regions that exhibited a difference in ISC between the Intact and Scrambled conditions for each group separately. The effect of interest for these models was condition (i.e., Intact and Scrambled) while controlling for episode and crossed random effects. Consistent with previous studies (Hasson et al. 2008, Lerner et al. 2011, Chen et al. 2016b), the Adult group exhibited greater ISC for the Intact compared to the Scrambled condition within the bilateral temporoparietal junction (TPJ), intraparietal sulci, supramarginal gyri, precuneus, middle inferior temporal gyri, and dorsal medial prefrontal cortex (dmPFC) (nodewise p<0.01, cluster extent 112 mm^2^, FWE p < 0.05)(Figure 4a). We also found that regions in the extrastriate cortex and superior temporal gyrus showed greater ISC in the Scrambled condition. The adult whole-brain results were corroborated through our ROI analyses in which, controlling for episode and crossed random effects, we detected significantly greater ISC during the intact episode within the dmPFC (β_condition_=0.029, 95% CI [0.021, 0.038], t(46)=4.74, p<0.001), TPJ (β_condition_=0.014, 95% CI [0.007, 0.021], t(46)=2.66, p<0.05) and precuneus (β_condition_=0.052, 95% CI [0.045, 0.060], t(46)=9.57, p<0.001). We also observed greater ISC in the scrambled compared to the intact condition in the auditory ROI (β_condition_=-0.011, 95% CI [-0.015, −0.006], t(46)=−3.39, p<0.01). Further, the Child group showed a qualitatively similar pattern compared to the Adult group (Figure 4b), however, in the ROI analyses we only observed greater ISC in the intact condition within the precuneus (β_condition_=0.017, 95% CI [0.011, 0.023], t(60)=3.99, p<0.001) and not the dmPFC (β_condition_=0.003, 95% CI [-0.003, 0.010], t(60)=0.73, p=1.00) or TPJ (β_condition_=0.002, 95% CI [-0.003, 0.007], t(60)=0.58, p=1.00). We also observed a similar trend of greater ISC during scrambled in the auditory region in the child group (β_condition_=−0.010, 95% CI [−0.014, −0.006], t(60)=−3.61, p<0.01). All p-values were corrected for the four multiple comparisons of each ROI using a Bonferroni correction.

**Figure 4.**
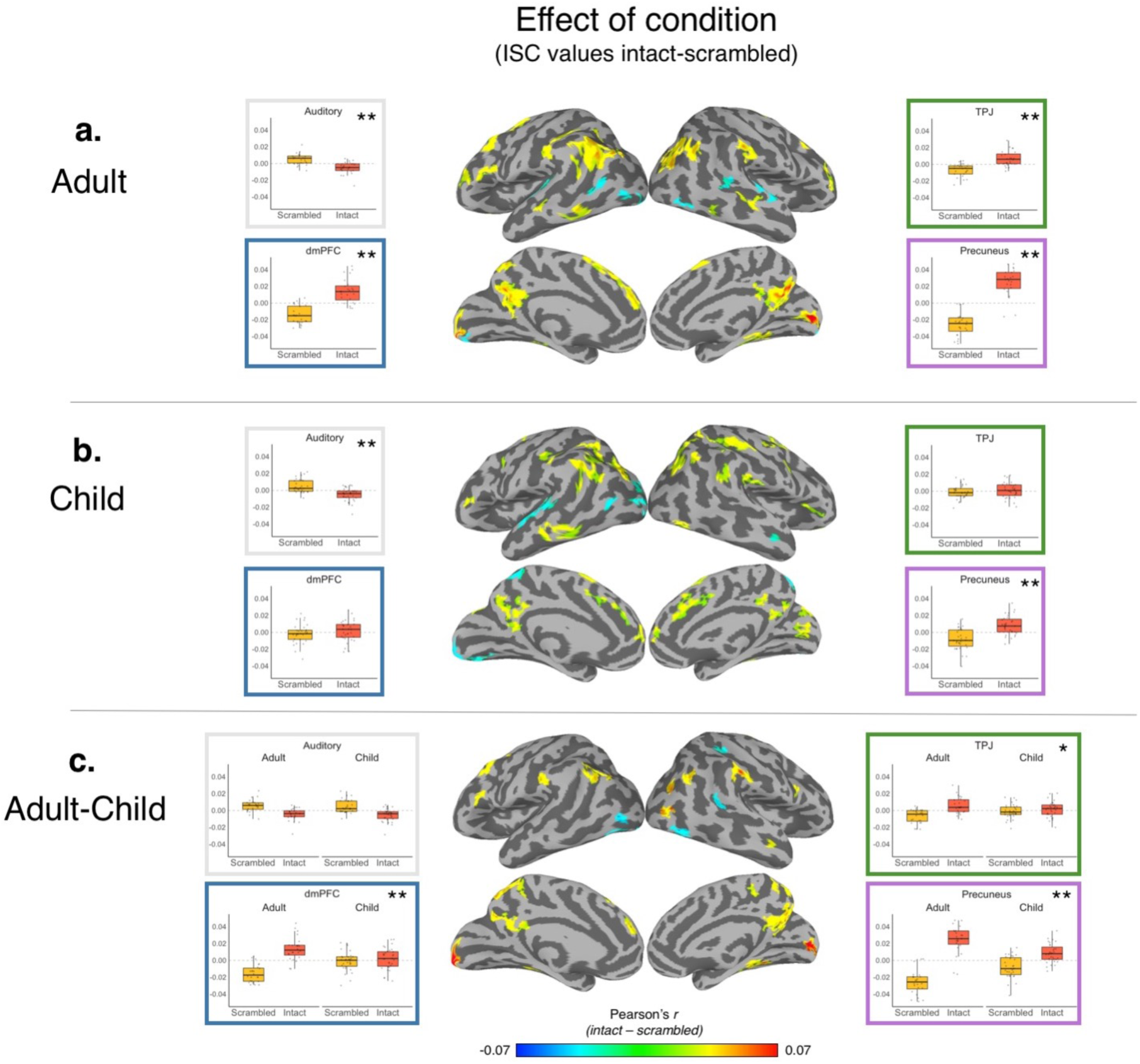
Functional specialization for long timescales. Intact versus scrambled ISC contrasts were created for the adult (a) and child (b) groups. The difference between the adult and child groups is shown in (c). In (a) and (b) hotter colors denote greater specialization for long timescale processing (e.g., intact-scrambled inter-subject correlation values) whereas in (c) hotter colors denote greater long timescale processing in the adult group (e.g., the effect of group on the difference between intact and scrambled). Whole brain maps are thresholded with a nodewise p < 0.01 with a cluster extent of 112 mm^2^ to achieve a FWE of p < 0.05. Each boxplot depicts the ISC values extracted from each region, controlling for episode and crossed random effects and then averaged for each participant. p values over the boxplots are Bonferroni corrected over the four ROI tests. * p<0.05, **p<0.001.

Finally, we examined which regions exhibited statistically greater selectivity for long timescales in the Adult compared to the Child group. Here all within-group ISC data for both groups were entered into the model. The effect of interest was the group by condition interaction while controlling for episode and crossed random effects. We found that the bilateral TPJ, supramarginal gyri, dorsal lateral prefrontal cortex (dlPFC), dorsal medial prefrontal cortex (dmPFC), inferior frontal gyri (iFG), precuneus, inferior temporal, and extrastriate cortices exhibited a group by condition interaction such that the difference between the Intact and Scrambled conditions was greater in the Adult compared to the Child group (nodewise p<0.01, cluster extent 112 mm^2^, FWE p < 0.05)(Figure 4c). In addition, we also observed statistically significant group by condition interaction effects witin the dmPFC (β_interaction_=0.027, 95% CI [0.016, 0.037], t(108)=3.54, p<0.01) and precuneus (β_interaction_=0.034, 95% CI [0.025, 0.044], t(108)=4.93, p<0.001) ROIs but not the auditory ROI (β_interaction_=0.000, 95% CI [-0.006, 0.006], t(108)=-0.06, p=1.00). We observed a marginal interaction effect within the TPJ (β_interaction_=0.010, 95% CI [0.002, 0.019], t(108)=1.64, p=0.11) which did not survive Bonferroni correction (p=0.42).

### Long timescales are important for social-cognitive comprehension in middle childhood

To examine the relationship between long timescale processing and social-cognitive comprehension, we first ran a mixed-effects model that used factors of condition (intact / scrambled), question type (mental / non-mental) and a condition by type interaction to predict episode comprehension, while also controlling for age (in the case of the child group) and episode. In the child group, we found a condition by question type interaction, such that the difference between mental and non-mental comprehension was greater in the scrambled compared to the intact episode (β=13.1, 95% CI [0.04,22.0], t(90)=2.94, p<0.01 – reported effects are in units of percent change in comprehension) (Figure 5a). Thus, consistent with our predictions, having information presented at long (compared to short) timescales was more important for mental state compared to non-mental state comprehension in the child group. We also detected a main effect of condition such that comprehension scores for the scrambled episode were lower than the intact episode (β=-19.5, 95% CI [-25.6,-13.1], t(89)=-6.20, p<0.001). We did not detect a main effect of question type (i.e., mental vs. non-mental), however numerically participants scored better on the non-mental compared to the mental questions (β=0.06, 95% CI [0.0,12.3], t(89)=1.92, p=0.06). In addition, a main effect of age was also detected such that episode comprehension increased as a function of child age (in months) (β=5.5, 95% CI [2.9,8.1], t(29)=4.21, p<0.001) (Figure 5b). In contrast, the adult group did not exhibit a condition by question type interaction (β=2.3, 95% CI [-7.3,12.0], t(68)=0.48, p=0.63). We detected a main effect of condition such that comprehension scores for the scrambled episode were lower than the intact episode (β=-17.6, 95% CI [-24.4,-10.7], t(68)=-5.10, p<0.001). We also detected a main effect of question type such that scores on the non-mental questions were higher than the mental questions) (β=16.4, 95% CI [9.5,23.4], t(68)=4.76, p<0.001) (Figure 5c).

**Figure 5.**
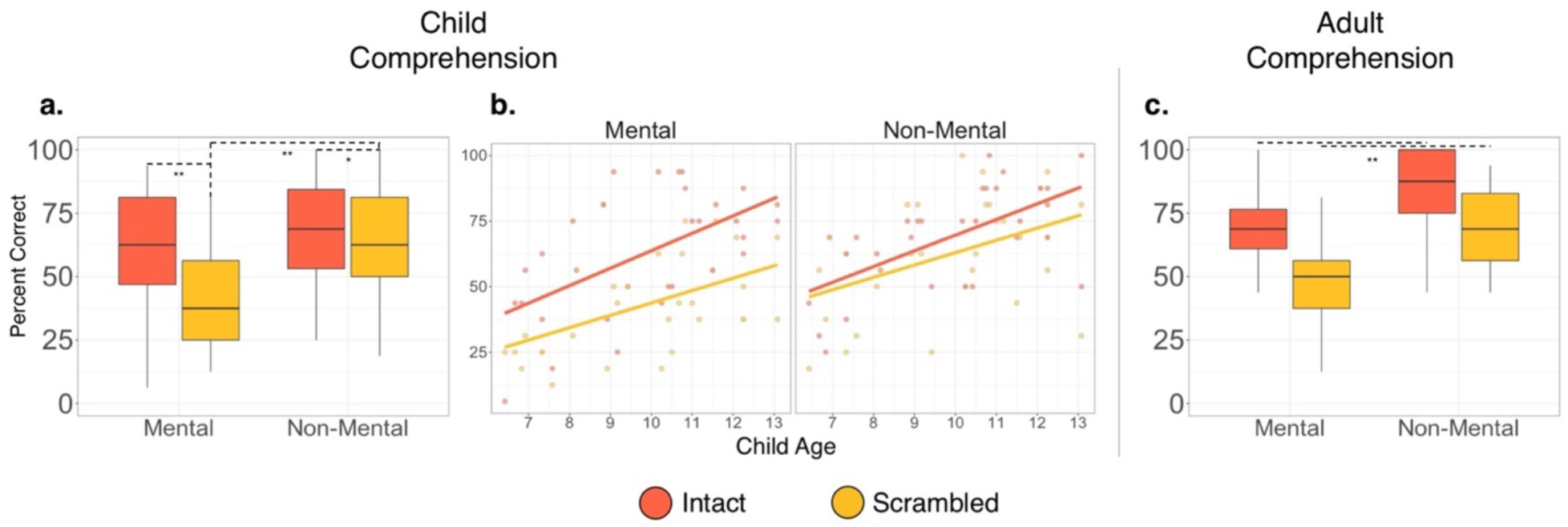
Episode comprehension. Comprehension is presented by question type (mental and non-mental) and condition (intact and scrambled) for the child group (a), as a function of child age (b), and for the adult group (c). *p<0.01, **p<0.001.

### Individual differences in neural maturity

We next examined how long timescale child-to-adult ISC varied as a function of individual differences in child age and mental state comprehension. Here our effect of interest was a covariate by condition interaction such that, we expected the covariates to be positively related to child-to-adult ISC within the intact episode more so than the scrambled episode. Controlling for episode and child head motion, we found that child age predicted child-to-adult whole-brain ISC in the left TPJ, right supramarginal gyrus, and bilateral extrastriate regions more so in the intact compared to the scrambled episode (nodewise p<0.01, cluster extent 112 mm^2^, FWE p < 0.05)(Figure 6a). In the ROI analyses, we detected a statistically significant age by condition interaction within the dmPFC (β_interaction_=0.004, 95% CI [0.002, 0.007], F(1,1390)=11.21, p<0.001) and the precuneus (β_interaction_=0.004, 95% CI [0.002, 0.007], F(1,1390)=11.62, p<0.001), such that child-to-adult ISC increased with age during the intact and not the scrambled episode, but not the TPJ (β_interaction_=0.000, 95% CI [-0.002, 0.003], F(1,1390)=0.46, p=0.50), or the auditory regions (β_interaction_=0.000, 95% CI [-0.001, 0.002], F(1,1390)=1.27, p=0.26).

**Figure 6.**
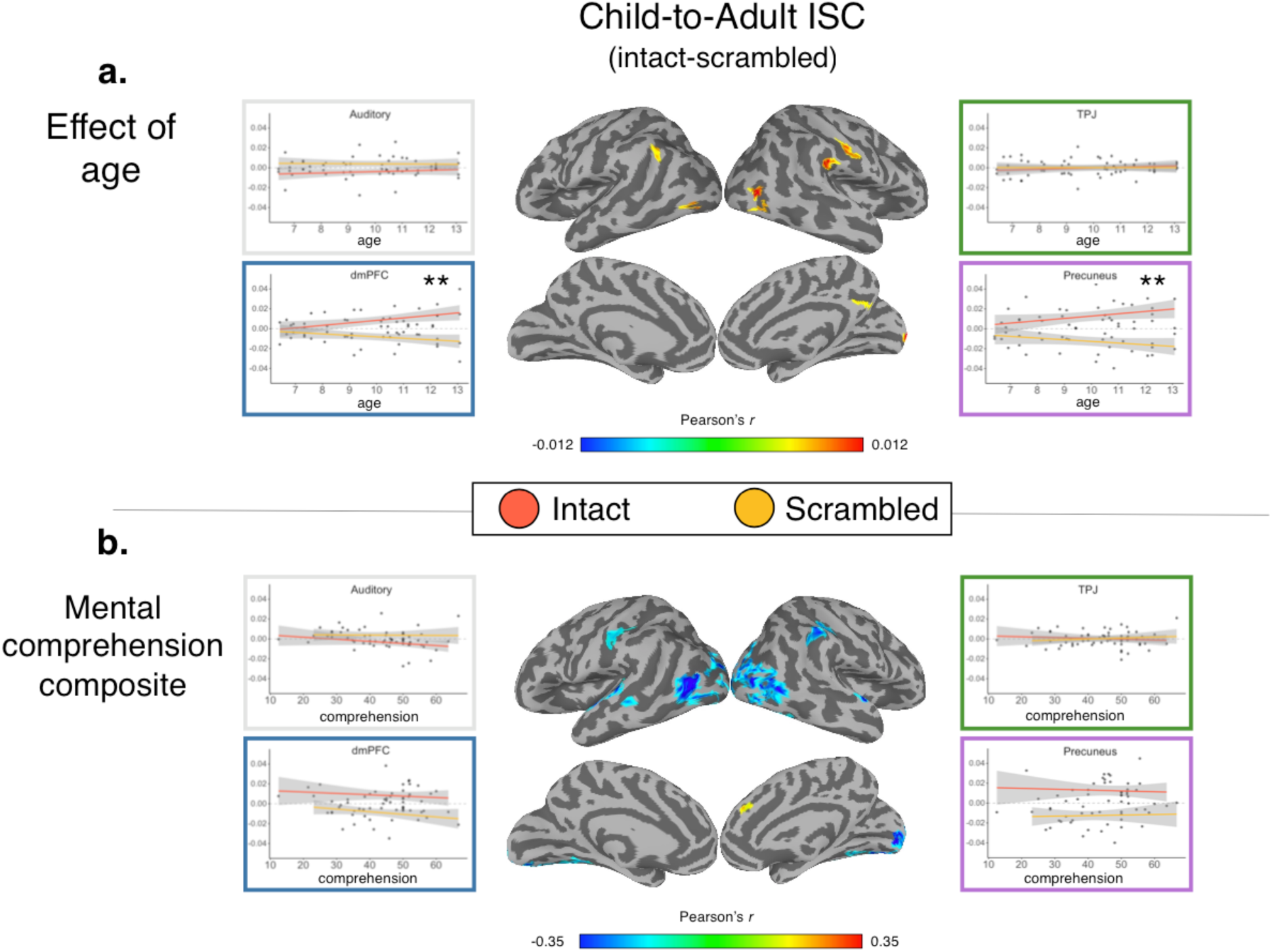
Child-to-adult ISC. The relationship between child-to-adult ISC and individual differences in child age (a) and mental state comprehension (b) were examined. The whole-brain statistical maps reflect an interaction between condition (intact-scrambled) and the covariate. Scatterplots show the relationship between the covariate and each condition. **p<0.001.

For comprehension, we used a composite to index mental state comprehension beyond general comprehension (see Methods) as our covariate. Controlling for episode, child head motion, and child age, we found that mental state comprehension predicted child-to-adult ISC in the right dmPFC more so in the intact compared to the scrambled episode. Further, we found that child-to-adult ISC in the bilateral extrastriate cortices as well as somatosensory cortex, inferior parietal lobe, and secondary auditory cortices was related to general episode comprehension (nodewise p<0.01, cluster extent 112 mm^2^, FWE p < 0.05) (Figure 6b). In the ROI analyses, we did not detect a statistically significant mental comprehension by condition interaction in any of the four ROIs (dmPFC: β_interaction_=0.000, 95% CI [0.000, 0.001], F(1,1409)=0.23, p=0.63; TPJ: β_interaction_=0.000, 95% CI [-0.001, 0.000], F(1,1409)=1.72, p=0.19; precuneus: β_interaction_=0.000, 95% CI [-0.001, 0.002], F(1,1409)=1.43, p=0.23, auditory: β_interaction_=0.000, 95% CI [-0.001, 0.000], F(1,1409)=3.37, p=0.07). See Supplementary Figure 6 for unthresholded statistical maps. Finally, while the aforementioned analyses focused on the effect of the mental comprehension composite, we also present results from each comprehension type and episode condition in Supplementary Figure 7.

## Discussion

In this first study to address cortical temporal hierarchy in children, we found evidence that children (6-13 year-old) exhibit less functional specialization for long timescale processing compared to adults. Further, within the TPJ – a key region for social cognition and long timescale processing – children showed reduced power in low-frequency spectra than adults. Behaviorally in the child group, we found that information presented over longer timescales (i.e., the intact episode) leads to better comprehension for mental state information than information presented over relatively shorter timescales. Further, we found age-related differences in child-to-adult ISC within the dmPFC and precuneus and that more ‘adult-like’ response in the dmPFC during long timescale processing is related to better social-cognitive comprehension, suggesting a role of this region in timescale processing specific to mental state comprehension. To our knowledge, this is the first study to examine developmental differences in cortical temporal hierarchy and the role of long timescale processing in social-cognitive processing in middle childhood.

Long timescale processing is thought to reflect the influence of information over multiple minutes or longer (Hasson et al. 2008, Hasson et al. 2015). Here we replicated previous low-frequency power and ISC findings within the same participants, such that that regions topographically similar to the DMN (e.g., TPJ, precuneus, dmPFC) are implicated in long timescale processing in adults both through manipulation of the temporal order of videos and examination of the power spectral density during intact movie viewing. While ISC was higher within long timescale regions for intact than scrambled videos the reverse was seen for the control region of auditory cortex. Auditory cortex is a short timescale region, generally sensitive to stimuli on the order of seconds or slightly longer (e.g., words) (Lerner et al. 2011). Thus, while this finding was not predicted it also does not run counter to our hypothesis that this region would not demonstrate greater low-frequency power during intact conditions. Our study is an extension of previous work (Hasson et al. 2008, Lerner et al. 2011, Chen et al. 2016b), such that previous studies presented participants with the same content for each timescale condition, whereas we presented participants with different content for the scrambled and intact conditions. This enabled us to examine effects of scrambling on ISC and comprehension without the confound of repeated content exposure between conditions.

While the child group showed a qualitatively similar pattern of regions associated with long timescale processing as the adult group, we found stronger and more focal neural synchrony in the adults compared to children within the regions of the DMN (e.g., bilateral TPJ, precuneus, dmPFC, iFG), and visual/attentional processing (e.g., supramarginal gyri, extrastriate cortices) during the intact compared to the scrambled episode. In addition, using ROIs defined from a meta-analysis of theory of mind processing in adults, (Schurz et al. 2014), we found a similar pattern of greater functional specialization for long timescales in adults compared to the children within the TPJ, dmPFC, and precuneus. Further, this finding of greater long timescale specialization in adults compared to children was consistent with our finding of greater low-frequency power in adults compared to children within the TPJ ROI. Together, our results suggest that functional specialization of long timescale processing during naturalistic viewing is immature in middle childhood. This immaturity could reflect less distinction between the intact and scrambled episodes at the group level or more idiosyncratic timescale processing that becomes more uniform with age. However, given the fact that we also see group differences in low-frequency power, idiosyncratic long timescale processing may be unlikely since the power analysis is not affected by between-subject variability. These between-group ISC findings in the current study are consistent with our previous work that suggests greater neural synchrony within the DMN in adults compared to children (Moraczewski et al. 2018).

In addition to examining group differences between children and adults, we examined whether age-related differences were found within middle childhood (i.e., 6 to 13 years) using a measure of child-to-adult neural synchrony and examination of age-related differences in low-frequency power within the child group. Here we found mixed support for age-related change during middle childhood, with discrepancies arising between the whole-brain and ROI findings (Figure 6a). In the whole-brain maps, child-to-adult neural synchrony during the intact episode increased as a function of child age within the left TPJ, right supramarginal gyrus and precuneus, and extrastriate cortices more so than during the scrambled condition. However, in the ROIs only the precuneus showed a similar condition by age interaction in both ROI and whole-brain analyses. Within the dmPFC ROI, we found a significant condition by age interaction that was not seen in the whole-brain statistical map. This discrepancy is likely due to the averaging of nodes for the ROI analyses. Indeed, in the unthresholded map in Supplementary Figure 6, we see positive, but not statistically significant, interactions within the bilateral dmPFC. Within the TPJ ROI, we did not see a significant effect of age but we did see a significant effect in a left TPJ cluster within the whole-brain analysis. This discrepancy is not due simply to averaging the TPJ ROI because we did not see an effect in either LTPJ or RTPJ ROIs independently (Supplementary Figure 4a). What likely does account for the discrepancy between the whole-brain and ROI TPJ analyses is the different spatial locations of the LTPJs. The ROI TPJ was defined based on a meta-analysis of theory of mind processing in adults (Schurz et al., 2014). Given the functional heterogeneity within the TPJ (Igelström et al. 2015), one possibility is that the TPJ showing whole-brain age by condition effects in the current study is not a region associated with theory of mind processing. An alternate possibility is that both regions are associated with theory of mind processes but do not both demonstrate age-related changes. Several lines of data support this latter interpretation. First, the peak effect within the left TPJ (MNI: −54, −57, 32) corresponds to associations of ‘theory of mind’, ‘mind’, and ‘default’ from the neurosynth meta-analysis (Yarkoni et al. 2011). Second, our whole-brain peak LTPJ region is very similar to that found in a task-based fMRI study directly probing age-related change in selectivity for mental state reasoning in similar aged children (MNI: −48, −60, 30) (Gweon et al. 2012) and is similar to the region that displayed age-related changes in our previous study (Moraczewski et al. 2018). Finally, the TPJ ROI did not show age-related change in low-frequency power within the child group. Thus, the LTPJ identified in the whole-brain may be a region that changes with age in middle childhood but the LTPJ region identified in the Schurz et al., meta-analysis does not. Future work could include functional localizers to better define functional subregions of the TPJ to examine how those are related to the development of long timescale processing.

An immature cortical temporal hierarchy in children compared to adults could have important implications for the development of neural and cognitive systems. Temporal hierarchy in the cortex is organized from short to long temporal receptive windows (TRWs) (Hasson et al. 2008), which is also thought to reflect a region’s location along a unimodal to transmodal gradient, respectively (Margulies et al. 2016, Huntenburg et al. 2018). Longer timescale processing may reflect recurrent connections between higher-order association cortices (e.g., DMN) that enable the ability to integrate newly arriving information while also maintaining an internal working model of one’s environment and cognitive state (Honey et al. 2012, Chaudhuri et al. 2015, Déli et al. 2017). Given the immaturity of the DMN during middle childhood (Fair et al. 2009, Supekar et al. 2010, de Bie et al. 2012, Muetzel et al. 2016), perhaps the development of long timescale processing reflects the refinement of these cortical gradients; however the current study cannot address these hypotheses. Future work should examine the developmental relationship between temporal hierarchy and cortical gradients at multiple scales and how cognitive abilities are affected by this system’s maturation. Further, one important caveat of the current study is that we did not record physiological measurements in the scanner (e.g., heartbeat and respiration). Previous work suggests that such measures have important and regionally-specific (e.g., TPJ) implications on the BOLD signal and future work examining developmental differences in timescale processing should address this limitation (Birn et al. 2006, Wu and Marinazzo 2016).

This study was the first to examine whether long timescale processing is related to social cognition. We found that children’s social-cognitive performance was dependent on information presented over longer timescales; whereas general memory was not affected by timescales to the same extent. Previous work shows that one’s ability to maintain context (e.g., holding onto information during a task) is specifically important for theory of mind abilities in terms of perceiving one’s social status (Rizzo and Killen 2018), planning moves during chess (Powell et al. 2017), and ingroup versus outgroup dynamics (Mulvey et al. 2016). Further, theory of mind is a complex ability that involves the coordination of multiple skills (e.g., face recognition, emotion processing, predicting behavior) (Schaafsma et al. 2015), many of which operate on diverse timescales. Thus, longer timescales may be needed to track, accumulate, and refine an accurate representation of mental states (Déli et al. 2017), which is essential to one’s ability to predict the social environment (Koster-Hale and Saxe 2013). It is important to note that it is not possible to disentangle whether the integration of information over long timescales is a domain-general neural property that affects social cognition or whether disruption to a social narrative causes information about a character’s mental states or actions to be lost and thus impairs social-cognitive performance. We argue that these are two sides of the same coin. The fact that many social-cognitive judgments require integration of prior information about another person suggests that the neural machinery supporting these computations must be able to operate over a long temporal receptive window (Redcay & Moraczewski, under review).

While long timescales were important for children’s social-cognitive performance, we did not detect a significant interaction effect between condition and question type in the adult group. Similar to the child group, mental comprehension was greater in the intact compared to scrambled episode; however, unlike the child group, comprehension on the non-mental questions was also affected by timescale. Specifically, the non-mental comprehension from the intact episode was higher than any other condition. Because our choice of stimuli (children’s show) and comprehension questions were designed to test comprehension in middle childhood specifically, the questions may have been too simple to disentangle the effects of long timescale on comprehension in adults. An alternate possibility is that presentation of information over long timescales is equally beneficial for social-cognitive and general memory in adults. The unique advantage of long timescales to social-cognitive comprehension in childhood may be most important for a developing social-cognitive system. Future work should examine interactions between question type and timescales in adults to address these possibilities. Further, given that questions were created based on an existing video, the stimuli were not perfectly matched on language and memory demands. A more precise matching between mental and nonmental comprehension questions (e.g., matching on social/nonsocial features and number of agents/actions) will be important for directly comparing effects between mental and non-mental questions.

To examine how ISC over long timescales is related to social-cognitive comprehension, we tested for relations between child-to-adult ISC and social-cognitive comprehension (while controlling for overall comprehension) in the intact compared to scrambled conditions. Here we found that child-to-adult ISC for the long timescale condition increased in the right dmPFC as a function of mental state comprehension beyond general comprehension. We post-hoc examined unthresholded maps relating child-to-adult ISC in intact and scrambled conditions to mental and nonmental comprehension in order to better understand these effects. When examining these patterns it appears that this effect is driven by differences in the relations between mental state comprehension and child-to-adult ISC in the intact compared to scrambled conditions, with a slight positive relation in intact and negative relation in scrambled conditions. The dmPFC is associated with social-cognitive abilities (Schurz et al. 2014), specifically the representation of socially-relevant information (e.g., personality traits) during task (Saxe and Powell 2006), spontaneous mentalizing (Castelli et al. 2000, Moessnang et al. 2016), viewing social interactions (Wagner et al. 2016), as well as long timescale processing (Lerner et al. 2011, Chen et al. 2016b). Previous work posits that response in this region is crucial for the ability to build individual representations of the history and personality traits of others (Mitchell et al. 2006), which facilitates one’s ability to predict what a social partner may do next (Koster-Hale and Saxe 2013). Thus, we speculate that successful performance on the mental state questions (compared to the non-mental questions) in the current study requires an accurate representation of each individual character. Further, the fact that the relationship between child-to-adult ISC and mental state comprehension was stronger during the intact compared to the scrambled episode suggests the importance of building the representations of others across time in this region. However, the relationship between mental state comprehension and child-to-adult ISC in the dmPFC was only observed in the whole-brain, but not the ROI, analysis. The ROI defined from a meta-analysis (Schurz et al. 2014) was centered on the left dmPFC, while the significant cluster in the whole-brain analysis was seen in the right hemisphere. While the left and right dmPFC are spatially proximal within 3D space, the bifurcation into left and right surfaces may have contributed to this discrepancy. Post-hoc examination in neurosynth (Yarkoni et al. 2011) of the peak coordinate within the right dmPFC cluster exhibiting the mental state comprehension by condition interaction (MNI: 11, 43, 33) suggest functional association terms of ‘mind tom’, ‘tom’, and ‘default network’, which provides further evidence of mental state processing within his region. However, future work should utilize a social-cognitive localizer to examine individual differences in mental state representation during naturalistic viewing.

In addition to a positive relation in the dmPFC between social-cognitive comprehension and child-to-adult ISC, we saw unexpected negative relations between child-to-adult ISC and the social-cognitive ratio comprehension score within clusters comprising ventral visual cortex, somatosensory cortex, and inferior parietal lobe. These are regions primarily comprising short and medium timescales (Lerner et al. 2011, Chen et al. 2016b). From examining the maps by condition and comprehension type it appears that this effect is largely driven by the difference in relations between nonmental comprehension with the intact videos (positive) and with the scrambled videos (negative). This effect was not predicted and requires further study to understand why these regions not typically associated with long timescales would show a stronger behavioral relation to intact compared to scrambled child-to-adult ISC. However, the stronger relation to nonmental comprehension within ventral visual, somatosensory, and inferior parietal regions is broadly consistent with research demonstrating greater activation of these regions when participants make judgments about physical, nonmental characteristics of a person, like bodily sensations (Jacoby et al. 2016). Future work is needed to clarify whether these processes similarly show greater disruption by long timescale processing as our current behavioral data in children do not support that claim.

In conclusion, our study is the first to investigate developmental differences in cortical temporal hierarchy, as well as the relationship between long timescale processing and social-cognitive abilities in middle childhood. We found that children exhibit immature long timescale processing within regions of the DMN. Further, during this age long timescales are more important for social-cognitive abilities beyond general memory. Finally, our data suggests that the dmPFC is important in the development of theory of mind, specifically in the representation of information about individuals over time.

## Acknowledgements

This work was supported by the National Science Foundation research traineeship program fellowship (award number: 1632976) awarded to D.M., the University of Maryland Ronald E. McNair Scholar Program awarded to J.N., University of Maryland funds, and National Institute of Mental Health grant (RO1MH112517) awarded to E.R.

The authors would like Aiste Cechaviciute, Sabine Huber, Alex Mangerian, Sydney Maniscalco, Hunter Rogoff, and Tova Rosenthal for assistance with data collection and data quality assessment and the Maryland Neuroimaging Center and staff for support with data collection.

