## Supplementary Information for "Cortical temporal hierarchy is immature in middle childhood"

#### Sample head motion

After our exclusion of motion outliers using mean frame displacement (FD) (see Methods in the manuscript), our final sample was 31 participants in the child group (mean(SD) FD = 0.16(0.04)) and 24 in the adult group (mean(SD) FD = 0.14(0.04)). Using a mixed-effects model to account for repeated measures, we did not detect a difference in mean FD between the adult and child groups ($\beta_{group}$ = 0.014, 95% CI [-0.007, 0.034], t(62)=1.30, p=0.20), nor did we detect a difference in condition ($\beta_{condition}$ = 0.004, 95% CI [-0.004, 0.013], t(53)=0.99, p=0.33) or a group by condition interaction ($\beta_{interaction}$ = 0.005, 95% CI [-0.006, 0.017], t(53)=0.93, p=0.36) (Supplementary Figure 1a). All units of mean FD are in millimeters.

In the child group, we detected a negative effect of age on mean FD such that older children has less head motion compared to younger children ($\beta_{age}$ = -0.007, 95% CI [-0.014, -0.001], t(36)=-2.30, p<0.05). We did not detect a difference between condition ($\beta_{condition}$ = 0.004, 95% CI [-0.041, 0.049], t(29)=0.16, p=0.87) or an age by condition interaction in the child group ($\beta_{interaction}$ = 0.001, 95% CI [-0.004, 0.005], t(29)=0.28, p=0.80) (Supplementary Figure 1b).

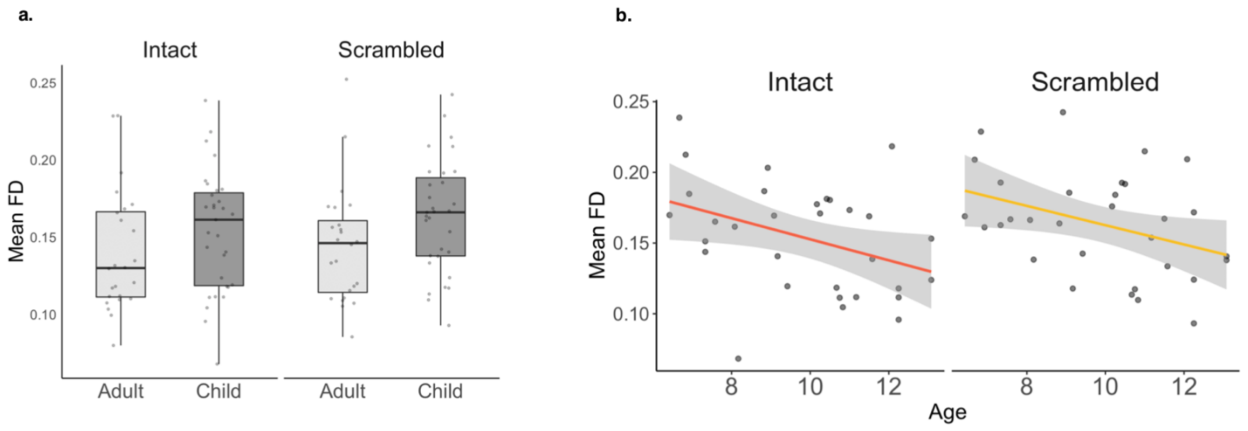


**Supplementary Figure 1 – Final sample head motion.** (a) Mean frame displacement by group and condition. (b) Mean frame displacement by condition and age in the child group. FD = frame displacement.

### Separate left and right TPJ analysis

In both the low-frequency power and inter-subject correlation (ISC) analyses in the manuscript, we represented the TPJ as an averaged left and right hemisphere ROI. This decision was made since, to our knowledge, no work has systematically examined hemispheric differences in long timescale processing. Here, we conduct the same analysis pipeline as in the manuscript while keeping the TPJ hemispheres separate.

### Low-frequency power in separate TPJ regions

Our results remain consistent with the averaged-TPJ results such that, while controlling for head motion, both the left and right TPJ regions exhibited greater low-frequency power in the adult compared to the child group (Left: $\beta_{group}$=0.07, 95% CI [0.03,0.11], t(52)=3.23, p<0.01; Right: $\beta_{group}$=0.05, 95% CI [0.01,0.10], t(52)=2.35, p<0.05) (Supplementary Figure 2a). In addition, as with the averaged-TPJ results, controlling for head motion, we did not observe a relationship between child age and proportion of low-frequency power (Left: $\beta_{age}$=0.01, 95% CI [-0.01,0.02], t(28)=1.61, p=0.51; Right: $\beta_{age}$=0.00, 95% CI [-0.02,0.01], t(28)=-0.47, p=1.00) (Supplementary Figure 2b).

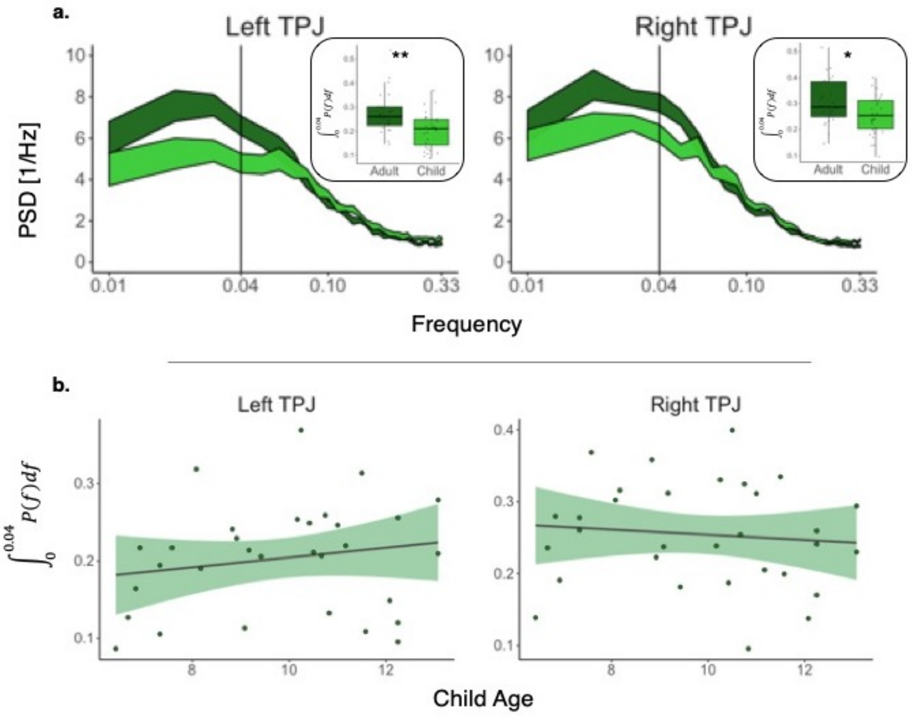


**Supplementary Figure 2 – Low-frequency power in separate TPJ regions.** (a) Power spectrum density (PSD) for the adult and child groups within the left and right TPJ regions. The PSD ribbons for each region depict the mean ± standard error. The black line denotes the 0.04 Hz cut-off used in the low-frequency statistical analysis. The boxplots denote the proportion of low-frequency power for each region. (b) Relationship between the proportion of low-frequency power and child age within the child group. * p<0.05, ** p<0.01, controlled for multiple comparisons.

**Functional specialization for long timescales in separate left and right hemisphere TPJ**

We next examined functional specialization for long timescales within the left and right TPJ regions separately. As in the main manuscript, we extracted average pairwise ISC values from each ROI and entered them into a mixed-effects model, controlling for episode and crossed random effects. In the adult group, we found that both the left and right TPJ regions exhibited greater ISC during the intact compared to the scrambled conditions, however, after correcting for the two comparisons using a Bonferroni correction, only the right TPJ remained significant. (Left: $\beta_{condition}$=0.014, 95% CI [0.003, 0.024], t(46)=1.75, p=0.17; Right: $\beta_{condition}$=0.014, 95% CI [0.006, 0.022], t(46)=2.40, $p<0.05$) (Supplemental Figure 3a).

In the child group, neither the left or right TPJ regions exhibited a significant difference in ISC between the intact and scrambled conditions (Left: $\beta_{condition}$=0.001, 95% CI [-0.006, 0.012], t(60)=0.45, p=1.00; Right: $\beta_{condition}$=0.001, 95% CI [-0.004, 0.007], t(60)=0.37, p=1.00) (Supplemental Figure 3b).

To examine the group by condition interaction, we entered all within-group pairwise ISC data into another mixed-effects model. Controlling for episode and crossed random effects, we did not detect a significant group by condition interaction in either the left or right TPJ regions (Left: $\beta_{interaction}$=0.000, 95% CI [-0.007, 0.022], t(108)=0.74, p=0.92; Right: $\beta_{interaction}$=0.013, 95% CI [0.003, 0.022], t(108)=1.91, p=0.12) (Supplementary Figure 3c).

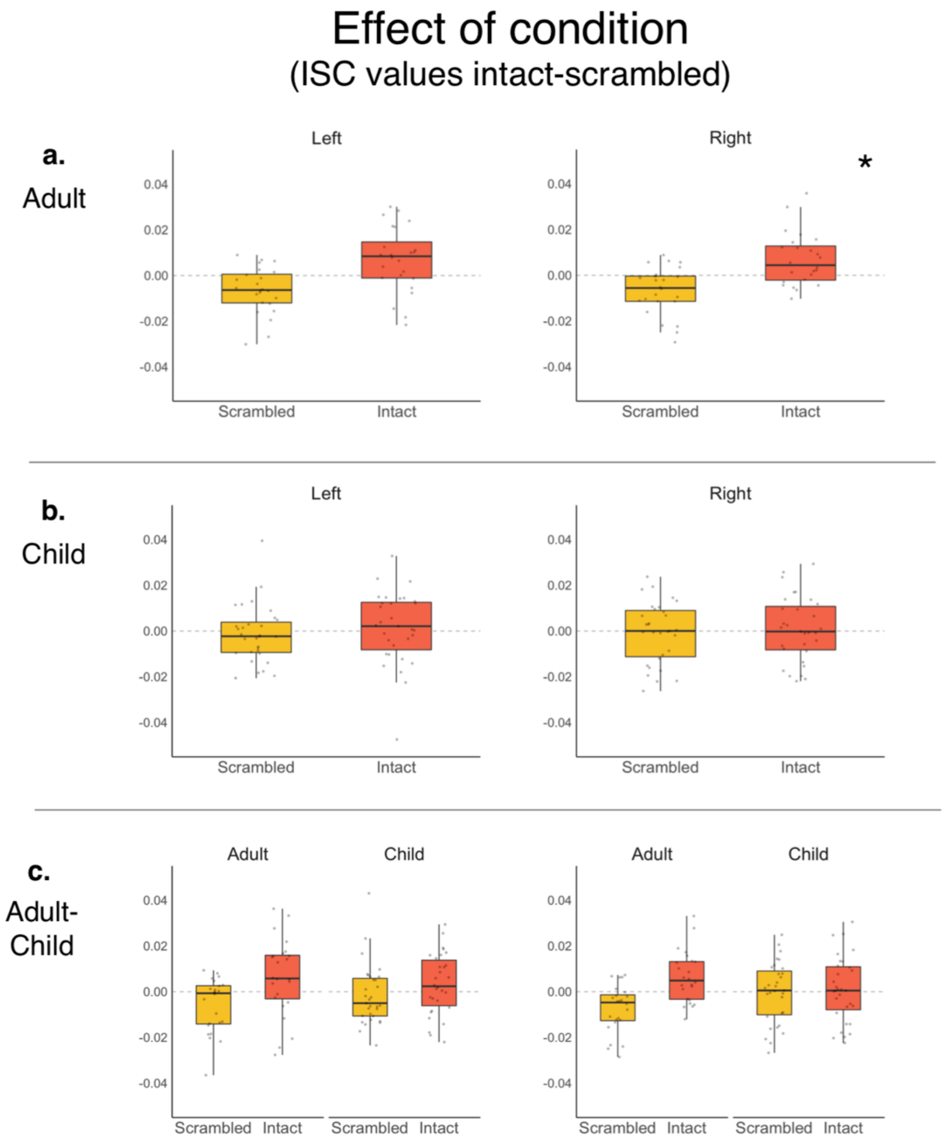


**Supplementary Figure 3 – Functional specialization for long timescales in separate TPJ regions.** Intact vs scrambled within-group ISCs are shown for the adult (a) and child (b) groups for the separate left and right TPJ regions. (c) Depicts the ISCs with both groups in the same model to examine a group by condition interaction. All ISC values on the y-axis residuals (after regressing out episode and crossed random effects) averaged over each participant. * p<0.05, controlled for two comparisons (left and right regions) using a Bonferroni correction.

### Individual differences in neural maturity in separate left and right hemisphere TPJ

Finally, as in the main manuscript, we examined the relationship between child-to-adult ISC and individual differences in age and mental state comprehension. The results from the separate TPJ hemispheres remain consistent with the average region presented in the manuscript. Controlling for episode and head motion, we did not detect an age by condition interaction on child-to-adult ISC in either hemisphere (Left: $\beta_{interaction}$=0.001, 95% CI [-0.002, 0.005], t(106)=0.99, p=0.65; Right: $\beta_{interaction}$=0.000, 95% CI [-0.002, 0.002], t(106)=-0.05, p=1.00) (Supplementary Figure 4a). Similarly, we did not detect a mental state composite comprehension by condition interaction on child-to-adult ISC (Left: $\beta_{interaction}$=-0.001, 95% CI [-0.001, 0.000], t(106)=-1.20, p=0.46; Right: $\beta_{interaction}$=0.000, 95% CI [-0.001, 0.000], t(106)=-0.27, p=1.00) (Supplementary Figure 4b). All p values were Bonferroni corrected for the two tests for each hemisphere.

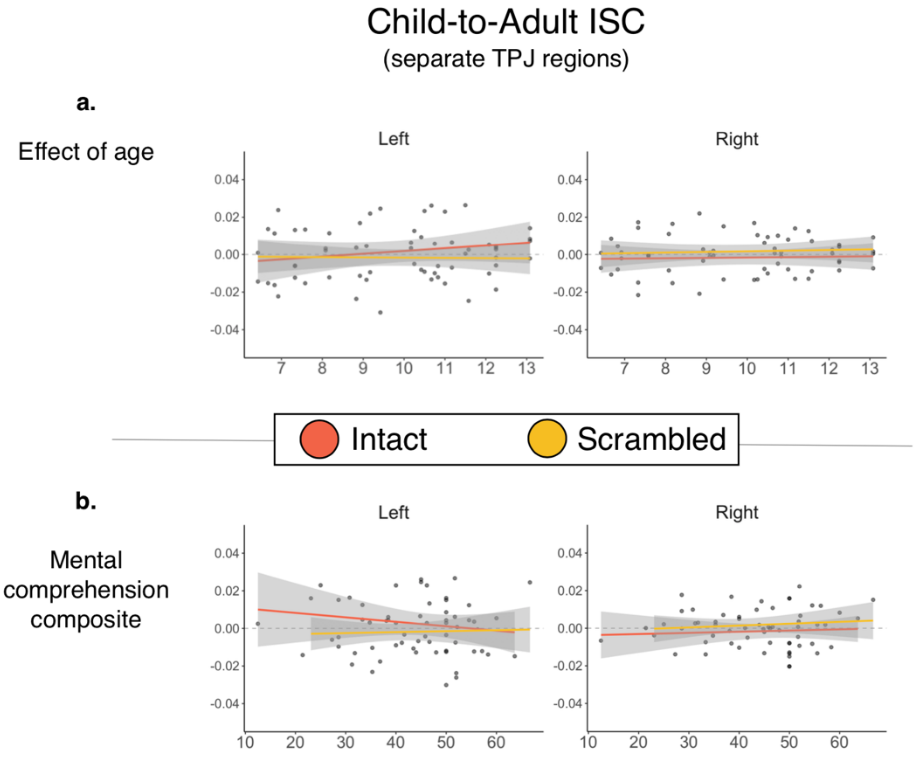


**Supplementary Figure 4 – Child-to-adult ISC in separate TPJ regions.** The relationship between child-to-adult ISC and individual differences in child age (a) and mental state comprehension (b) were examined.

### Effect of episode on specialization for long timescales

While our main question-of-interest revolved around neural specialization for long timescale processing regardless of episode content (e.g., which episode was scrambled versus intact), participants viewed different content for each of the intact and scrambled conditions. Thus, in all our inter-subject correlation (ISC) analysis we control for the effect of episode (e.g., “The Monkey Bars” (MB) or “The Body Bus” (BB)) by adding a fixed effect of episode into each regression model examining ISC for the intact and scrambled episodes (as well as their contrast). While we did not have a priori hypotheses about the effect of content (e.g., MB versus BB), we wanted to control for possible variation between episodes. Below we present whole-brain maps that show how specialization for long timescales (e.g., intact minus scrambled) differs between episode content (Supplementary Figure 5).

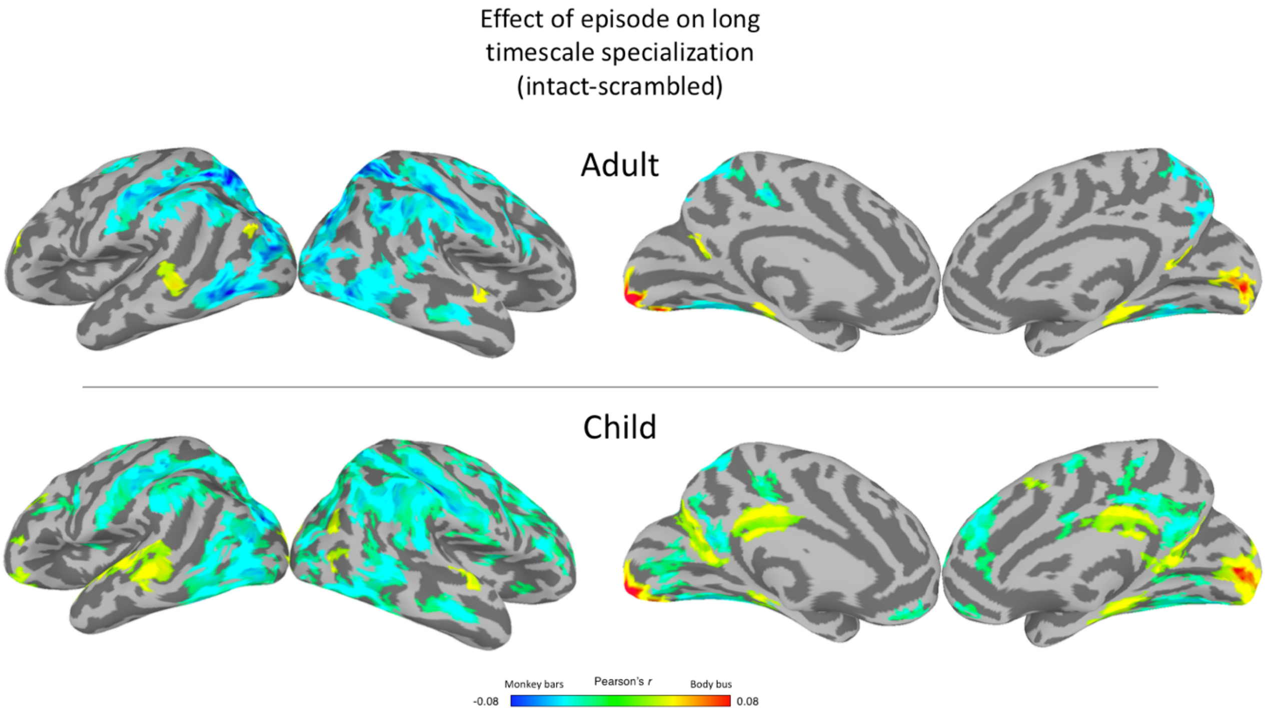


**Supplementary Figure 5 –** *Effect of episode*. The effect of episode (MB versus BB) on intact versus scrambled ISC for the adult (a) and child (b) groups. Maps are thresholded with a nodewise $p < 0.01$ with a cluster extent of 112 $mm^2$ to achieve a FWE of $p < 0.05$.

### Individual differences in neural maturity (unthresholded)

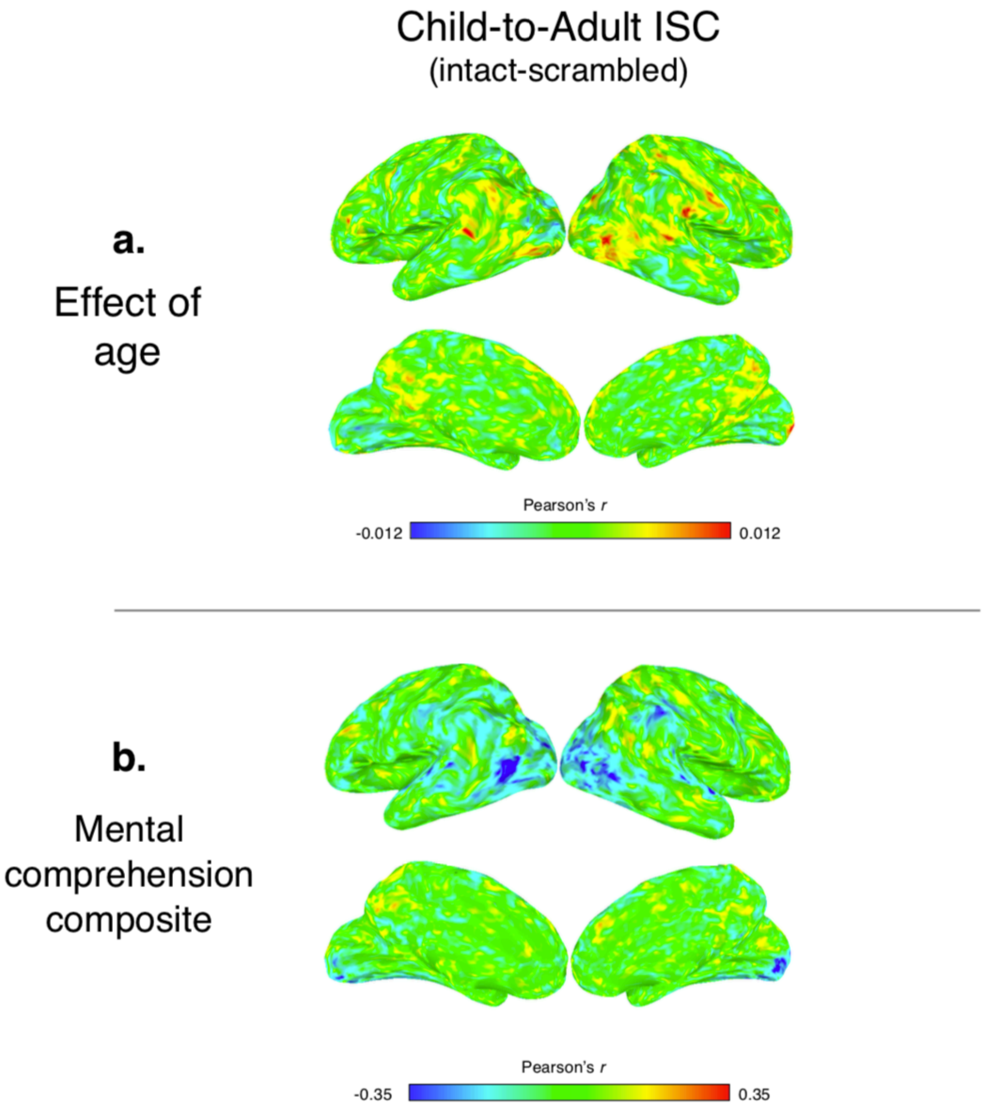


**Supplementary Figure 6 – Child-to-adult ISC.** The relationship between child-to-adult ISC and individual differences in child age (a) and mental state comprehension (b) were examined. The results here are the same as in Figure 6 of the manuscript, however these are unthresholded.

### Effect of comprehension type on child-to-adult neural synchrony

In the manuscript, our analyses on the effect of episode comprehension on child-to-adult neural synchrony focused on a mental state composite in which we controlled for overall comprehension (mental / mental + nonmental). Here we present unthresholded whole-brain maps to show the effect of each comprehension type (mental and nonmental) on child-to-adult neural synchrony within the intact and scrambled episodes separately (Supplementary Figure 6).

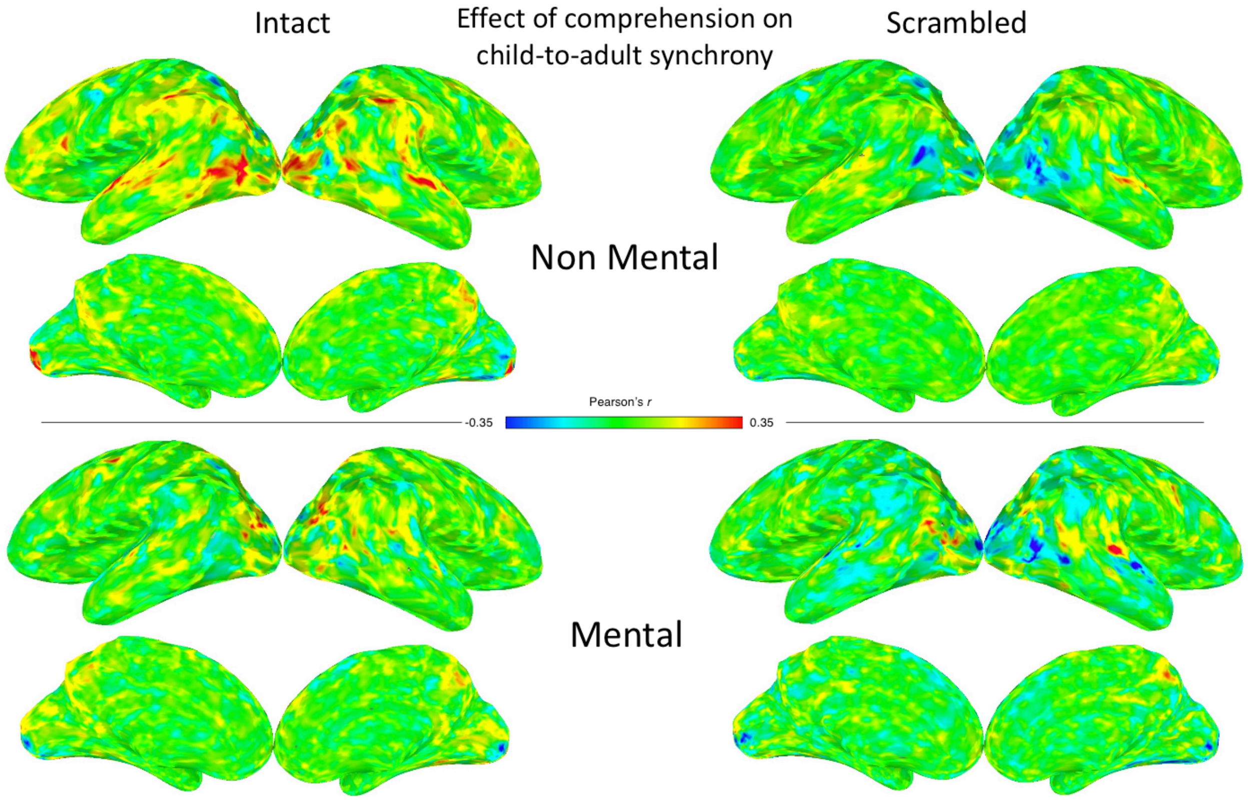


**Supplementary Figure 7 –** *Effect of comprehension type on child-to-adult neural synchrony.*

**Appendix 1: Comprehension questions**

**Season 1, Episode 4: The Monkey Bars**

**Mental questions**

1. Does Atticus think that Tamara thinks she owns the monkey bars?

| Score | 1 | 0.5 | 0 |
|---|---|---|---|
| Answer | Yes | N/A | No |

2. Why does Melanie think that she can't hang all day on the money bars?

| Score | 1 | 0.5 | 0 |
|---|---|---|---|
| Answer | She has swim practice after school | N/A | No mention of swim practice after school |

3. What does Battie think Tamara looks like when she is interrupted by Melanie?

| Score | 1 | 0.5 | 0 |
|---|---|---|---|
| Answer | Mean dog | Angry or upset | No mention of dog, angry, or upset |

4. Why is Battie glad that it wasn’t him who was hanging from the monkey bars?

| Score | 1 | 0.5 | 0 |
|---|---|---|---|
| Answer | He has weak arms and/or he had stinky breath because of something he ate last night | He didn’t want to fall off | No mention of weak arms, stinky breath, or fear of falling |

5. What does Tamara think that Melanie is looking for in her pocket?

| Score | 1 | 0.5 | 0 |
|---|---|---|---|
| Answer | Her handkerchief (or hanky) | N/A | No mention of handkerchief or hanky |

6. Why does Melanie think her teacher is in a hurry to leave?

| Score | 1 | 0.5 | 0 |
|---|---|---|---|
| Answer | To get another sausage | No mention that she is going back for seconds | No mention of sausages |

7. Why did Debra Jo ask the kids to calculate their own change?

| Score | 1 | 0.5 | 0 |
|---|---|---|---|
| Answer | To provide them with a learning experience | N/A | No mention of the intention to teach |

8. Why did Melanie not want to remind Tamara about the sausage sizzle?

| Score | 1 | 0.5 | 0 |
|---|---|---|---|
| Answer | She didn't want Tamara to get a sausage first | N/A | No mention of the worry about Tamara getting the sausage first |

**Non-Mental questions**

1. What does Rory eat for breakfast?

| Score | 1 | 0.5 | 0 |
|---|---|---|---|
| Answer | Cereal | N/A | No mention of cereal |

2. What is 'two-sies?

| Score | 1 | 0.5 | 0 |
|---|---|---|---|
| Answer | Skipping every other monkey bar | Mention monkey bar but not skipping bars | No mention of monkey bars or skipping bars |

3. What fruit is Tamara eating?

| Score | 1 | 0.5 | 0 |
|---|---|---|---|
| Answer | Banana | N/A | No mention of banana |

4. What sport is Atticus and Rory playing?

| Score | 1 | 0.5 | 0 |
|---|---|---|---|
| Answer | Soccer | N/A | No mention of soccer |

5. Why does Melanie have money?

| Score | 1 | 0.5 | 0 |
|---|---|---|---|
| Answer | For the sausage sizzle | To buy lunch | No mention of sausages or buying lunch |

6. Did Melanie spend her money on sausages?

| Score | 1 | 0.5 | 0 |
|---|---|---|---|
| Answer | No | N/A | Yes |

7. What was Melanie folding out of paper?

| Score | 1 | 0.5 | 0 |
|---|---|---|---|
| Answer | Fortune teller, cootie catcher, or proper participant hand motion | N/A | No mention of object name or hand motion |

8. What can Tamara do while she is hanging upside down on the monkey bars?

| Score | 1 | 0.5 | 0 |
|---|---|---|---|
| Answer | Mention krumps (the dance) | Dance, swing, or making the motion of the dance but not the name | No mention of krump, dancing, swinging, or not proper motion |

**Season 1, Episode 22: The Body Bus**

**Mental questions**

1. Why does Rory eat Melanie's sugar snap peas?

| Score | 1 | 0.5 | 0 |
|---|---|---|---|
| Answer | He wants to be healthy or he doesn't want to get sick | N/A | No mention of health or not getting sick |

2. Why does Atticus ask for a toothbrush?

| Score | 1 | 0.5 | 0 |
|---|---|---|---|
| Answer | He thinks there is a dentist at school that day | He wants to brush his teeth and/or because he ate candy | No mention of dentist, brushing teeth, or candy |

3. Why does Melanie think Debra Jo walks away from Rory's questions?

| Score | 1 | 0.5 | 0 |
|---|---|---|---|
| Answer | She thinks Debra Jo is upset | N/A | No mention of upset |

4. Why does the teacher think Debra Jo wants to call her mom?

| Score | 1 | 0.5 | 0 |
|---|---|---|---|
| Answer | She forgot to make her bed | N/A | No mention of needing to make bed |

5. Does Atticus think Rory saw a documentary about robots?

| Score | 1 | 0.5 | 0 |
|---|---|---|---|
| Answer | No | N/A | Yes |

6. Why did Battie come over to the tree?

| Score | 1 | 0.5 | 0 |
|---|---|---|---|
| Answer | To hide, be alone, read, and/or read to the birds | To climb the tree | No mention of climbing the tree, hide, be alone, or read |

7. Why did Melanie and Tamara go over to the tree?

| Score | 1 | 0.5 | 0 |
|---|---|---|---|
| Answer | To eavesdrop on the conversation between Debra Jo and the teacher | To go see what Debra Jo was up to | No mention of Debra Jo or her conversation with the teacher |

8. Why did Rory say that he combs his hair for longer than three minutes?

| Score | 1 | 0.5 | 0 |
|---|---|---|---|
| Answer | He didn't want to seem odd to Melanie and Tamara | N/A | No mention of him saying that because of social pressure |

**Non-Mental questions**

1. What color was the truck?

| Score | 1 | 0.5 | 0 |
|---|---|---|---|
| Answer | Blue, blue and white | N/A | No mention of blue |

2. Why were Battie and Atticus's hands dirty?

| Score | 1 | 0.5 | 0 |
|---|---|---|---|
| Answer | They were gardening, planting, or composting | They were digging in the dirt | No mention of gardening, planting, composting, or dirt |

3. Who was sitting on the bench with Rory and Tamara?

| Score | 1 | 0.5 | 0 |
|---|---|---|---|
| Answer | Melanie | N/A | No mention of Melanie |

4. What object was in the plastic tub in the classroom?

| Score | 1 | 0.5 | 0 |
|---|---|---|---|
| Answer | Cell phones | N/A | No mention of phones |

5. How did Debra Jo line the kids up?

| Score | 1 | 0.5 | 0 |
|---|---|---|---|
| Answer | Alphabetical order in boy-girl pairs | Using a megaphone | No mention of alphabetical, boy-girl pairs, or megaphone |

6. Where is Debra Jo when she calls her mom during class?

| Score | 1 | 0.5 | 0 |
|---|---|---|---|
| Answer | Under the teacher's desk | In the classroom, but no mention of under desk | No mention of classroom or desk |

7. Did Debra Jo have a splinter rash headache when she was in the nurses office?

| Score | 1 | 0.5 | 0 |
|---|---|---|---|
| Answer | No | N/A | Yes |

8. What did the Body Bus actually say?

| Score | 1 | 0.5 | 0 |
|---|---|---|---|
| Answer | Peabody Business Supplies | 2 of the 3 words | 1 or no correct words |
